# Discovery of the metalloenzyme IsmB revises a pathway for coprostanol formation by the human gut microbiome

**DOI:** 10.64898/2025.12.29.696961

**Authors:** Antonio Tinoco, Chenhao Li, Jai K. Khurana, Bryan Bañuelos Jara, Damian R. Plichta, Douglas J. Kenny, Ramnik J. Xavier, Emily P. Balskus

## Abstract

High levels of circulating cholesterol are associated with human cardiovascular diseases and an altered gut microbiome. Still, major gaps exist in our understanding of the interactions of cholesterol with gut microbes. The reductive transformation of cholesterol to the poorly absorbed sterol coprostanol by human gut bacteria has long been known, but the genetic and biochemical basis for this activity is only partially elucidated. Here, we discover and characterize a gut bacterial enzyme that catalyzes the reduction of cholestenone to coprostanone, the second step in the intestinal sterol metabolism (*ism*) pathway for coprostanol production. We identify a gene encoding a previously unknown 5β-reductase, IsmB, a new member of the Fe–S cluster flavoenzyme superfamily, in the coprostanol producing organism *Eubacterium coprostanoligenes*. Biochemical characterization of IsmB confirms it is an anaerobic Fe–S cluster flavoenzyme and reveals specificity for reduction of an unanticipated intermediate, 5-cholesten-3-one, to coprostanone, revising the *ism* pathway. We also identify and characterize homologs of IsmB encoded in uncultured human gut bacteria that also encode the previously identified *ism* pathway enzyme IsmA, further supporting the role of IsmB in coprostanol formation. Finally, analysis of human stool metagenomics and metabolomics datasets further confirms the relevance of IsmB in the human gut microbiome, and analyses of human serum metabolomics from Framingham Heart Study participants reveal negative correlation between serum cholesterol levels and the presence of IsmA/IsmB encoders in the gut. Together, these results show the utility of combining biochemistry and stool metagenomic analysis for gut microbial enzyme discovery, and suggests IsmA/IsmB-encoding gut bacteria carry potential benefits for cholesterol homeostasis and cardiovascular health.

## INTRODUCTION

Microbes residing in the human gut have vast metabolic capabilities that strongly influence health and disease, and these activities extend the biochemical reactions supported by host cells.^1,2^ The production of gut microbial metabolites varies significantly across human subjects, and specific microbial metabolites are strongly correlated with health outcomes.^3,4^ Obtaining a biochemical understanding of how gut microbes produce metabolites and the impacts these metabolites have on host physiology is needed to accurately detect, interrogate, and manipulate specific metabolic functions in gut microbial communities and experimentally link them to disease. This challenge is exemplified by efforts to elucidate gut microbial metabolism of intestinal cholesterol, one of the most abundant and well-studied human-associated metabolites.^5,6^ It is biosynthesized *de novo* in the liver and is a common animal-derived lipid in the human diet, serving as a building block for vitamins, bile acids, and hormones.^5–7^ Increased serum cholesterol in humans is a major risk factor for cardiovascular disease, which is the leading cause of death for most Western industrialized countries.^8–10^ The human gut microbiome has been proposed to be an underappreciated contributor to circulating cholesterol level in humans.^11–14^ Associations between the gut microbiome community composition and circulating cholesterol levels revealed that the prevalence and abundance of specific gut bacteria likely contribute to the modulation of total cholesterol levels in humans.^15^ Recent work by our groups demonstrated a negative association between the human gut *Eubacterium* species,^12^ *Oscillibacter* species and *Alistipes obesi*^13^ and plasma cholesterol levels in human subjects from the Framingham Heart Study^13^.

Meanwhile, positive correlations between *Flavonifractor plautii*, *Eggerthella lenta*, and *Blautia hansenii*, among others, and plasma cholesterol levels were also measured.^13^

Human gut bacteria have been reported to chemically transform intestinal cholesterol leading to altered cholesterol regulation in hosts. The human gut commensal *Bacteroides thetaiotaomicron* (*Bt*) encoding a 3’-phosphoadenosine-5’-phosphosulfate-dependent sulfotransferase, Bt0416, performs conjugative chemistry on cholesterol to produce cholesterol sulfate and may be implicated in microbiome-dependent regulation of host cholesterol levels.^14,16^ Monocolonization experiments in germ-free mice with *Bt* wild type (WT) and *Bt*Δ0416 revealed that *Bt*Δ0416-colonized mice displayed lower levels of whole-blood cholesterol sulfate than those associated with *Bt* WT, and also displayed lower serum cholesterol levels, indicating that the conversion of cholesterol to cholesterol sulfate by gut microbes impacts both cholesterol sulfate and cholesterol homeostasis.^14^ Our recent discovery that *Oscillibacter* spp. isolates from human stool encode homologs of IsmA and convert cholesterol to 4-cholestenone, cholesterol ⍺-D-glucoside and 24 (*R*,*S*)-hydroxycholesterol.^13^ Additionally, humans with *Oscillibacter* and *Eubacterium* species in their gut microbiomes were found to have lower stool and serum cholesterol levels. These studies collectively reveal the ability of human gut bacteria to transform intestinal cholesterol into metabolites, and these gut microbial metabolic activities and products have the potential to modulate host cholesterol levels.

Gut bacteria also metabolize intestinal cholesterol to the poorly-absorbed sterol metabolite, coprostanol (**Figure 1A**), which is eliminated in feces,^17,18^ leading to the proposal that this activity depletes cholesterol in the small intestine leading to cholesterol-lowering effects.^19,20^ The first publicly-available coprostanol-producing strain, *Eubacterium coprotanoligenes* ATCC 51222 (*E. cop*)^21^, was isolated from a hog-sewage lagoon and reported to metabolize cholesterol to coprostanol through an indirect reductive pathway involving the oxidation of cholesterol to 5-cholesten-3-one (5-cholestenone) and the isomerization of the Δ^5,6^ double bond to produce 4-cholesten-3-one (4-cholestenone) (**Figure S1A**).^22^ The Δ^4,5^ double bond in 4-cholestenone is then reduced to form the 5β-configured coprostanone followed by reduction of the ketone to generate coprostanol. This changes the conformation of the sterol skeleton from a ‘linear’ to a ‘bent’ geometry. A direct pathway for the reduction of cholesterol to coprostanol has also been proposed (**Figure S1A**) and was recently reported in *Limosilactobacillus fermentum* monocultures isolated from raw yak milk,^23–25^ expanding our understanding of the bacteria capable of performing this transformation.

**Figure 1.**
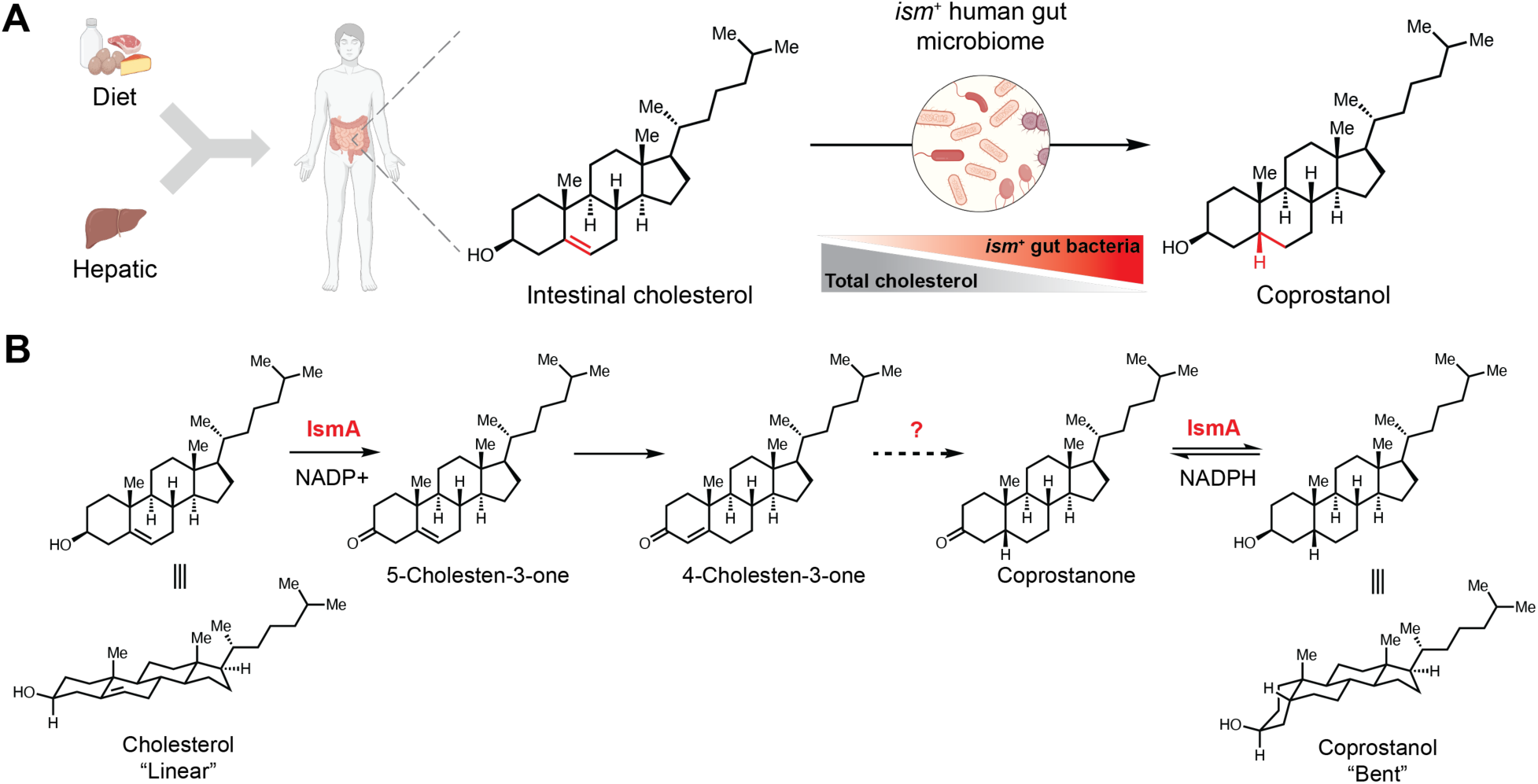
The human gut bacterial metabolism of intestinal cholesterol to the poorly absorbed sterol coprostanol is not fully understood. (A) The presence of *ism*^+^ species in a human gut microbiome correlates with lowered total serum cholesterol levels. (B) Previous proposed *ism* pathway involves the enzyme IsmA, which catalyzes the oxidation of cholesterol and isomerization of the C5–C6 double bond of 5-cholesten-3-one (5-cholestenone) to 4-cholesten-3-one (4-cholestenone) and the reduction of 5β-configured coprostanone to coprostanol. The gene(s) and enzyme(s) responsible for the reduction of 4-cholestenone to coprostanone was unknown.

The gut microbial genes and enzymes involved in the intestinal sterol metabolism (*ism*) pathway for the conversion of cholesterol to coprostanol were unknown until our discovery of the first enzyme in this process.^12^ We found that the intestinal sterol metabolism A gene (*ismA*) in the environmental isolate *E. cop* encodes a 3β hydroxysteroid dehydrogenase enzyme (IsmA) that catalyzes both the oxygen-independent oxidation and isomerization of cholesterol to 4-cholestenone and the reduction of coprostanone to coprostanol in a NADP^+^/NADPH-dependent manner (**Figure 1B**). Thus far, homologs of IsmA are found only in metagenomic species pangenomes (MSPs) assembled from fecal metagenomic sequencing data, and not in human gut-derived bacterial isolates. Human gut MSPs encoding IsmA homologs strongly correlate with coprostanol levels in stool metabolomes and with coprostanol production in human stool cultures. Finally, analyses of human clinical data revealed that participants with *ismA*^+^ gut microbiomes have lower stool cholesterol levels and lower total serum cholesterol levels compared to participants lacking *ismA*. Our work revealed, for this first time, that coprostanol-producing human gut bacteria may lower serum cholesterol levels, and thus have an impact on cardiovascular health.

Though the discovery of *ismA* advanced our understanding of the gut bacterial conversion of cholesterol to coprostanol, the gene(s) and enzyme(s) responsible for the second step in the previously proposed *ism* biosynthetic pathway, the stereoselective reduction of 4-cholestenone to coprostanone, remains unknown. Here, we use a chemically-guided genome mining approach to identify a gene and enzyme responsible for the second reductive step in this pathway in *E. cop*.

This enzyme, IsmB, is a flavin and iron–sulfur cluster-dependent 5β-reductase that catalyzes the key C–C double bond reduction to generate coprostanone. Unexpectedly, IsmB specifically accepts 5-cholestenone to produce 5β-configured coprostanone, revising the *ism* pathway. Lysate assays suggest that IsmB is constitutively expressed in *E. cop*. Homologs of IsmB are encoded in uncultivated human gut bacteria that also encode IsmA, and the presence of these genes in MSPs strongly correlates with coprostanol in fecal metabolomes, further supporting the role of IsmB in bacterial coprostanol production. Overall, our findings enhance our understanding of how gut bacteria convert cholesterol to coprostanol.

## RESULTS

### Identification of the enzyme responsible for the second step in microbial coprostanol production

We set out to discover the putative gut microbial enzyme responsible for converting 4-cholestenone to 5β-configured coprostanone. To do so, we took a chemically-guided genome mining approach to identify and prioritize candidate genes and enzymes with the catalytic capabilities needed to perform this reductive transformation in the coprostanol-producing bacterium *E. cop*. Specifically, we noticed parallels between this reaction and that catalyzed by the enzyme BaiCD encoded in the bile acid inducible (*bai*) operon found in gut commensal Clostridium strains.^26–28^ The *bai* pathway is responsible for the 7α-dehydroxylation of the primary bile acids cholic acid and chenodeoxycholic acid to produce deoxycholic acid and lithocholic acid, respectively.^26^ The BaiCD homolog in *Clostridium scindens* (*C. scindens*) is an oxygen-sensitive Fe–S cluster-dependent flavoenzyme and a member of the NADH- and flavin-dependent oxidoreductase superfamily^29,30^ that stereoselectively reduces 3-oxo-4,5-dehydro deoxycholic acid (DCA) to 5β-configured 3-oxo DCA^30^ (**Figure 2A**).

**Figure 2.**
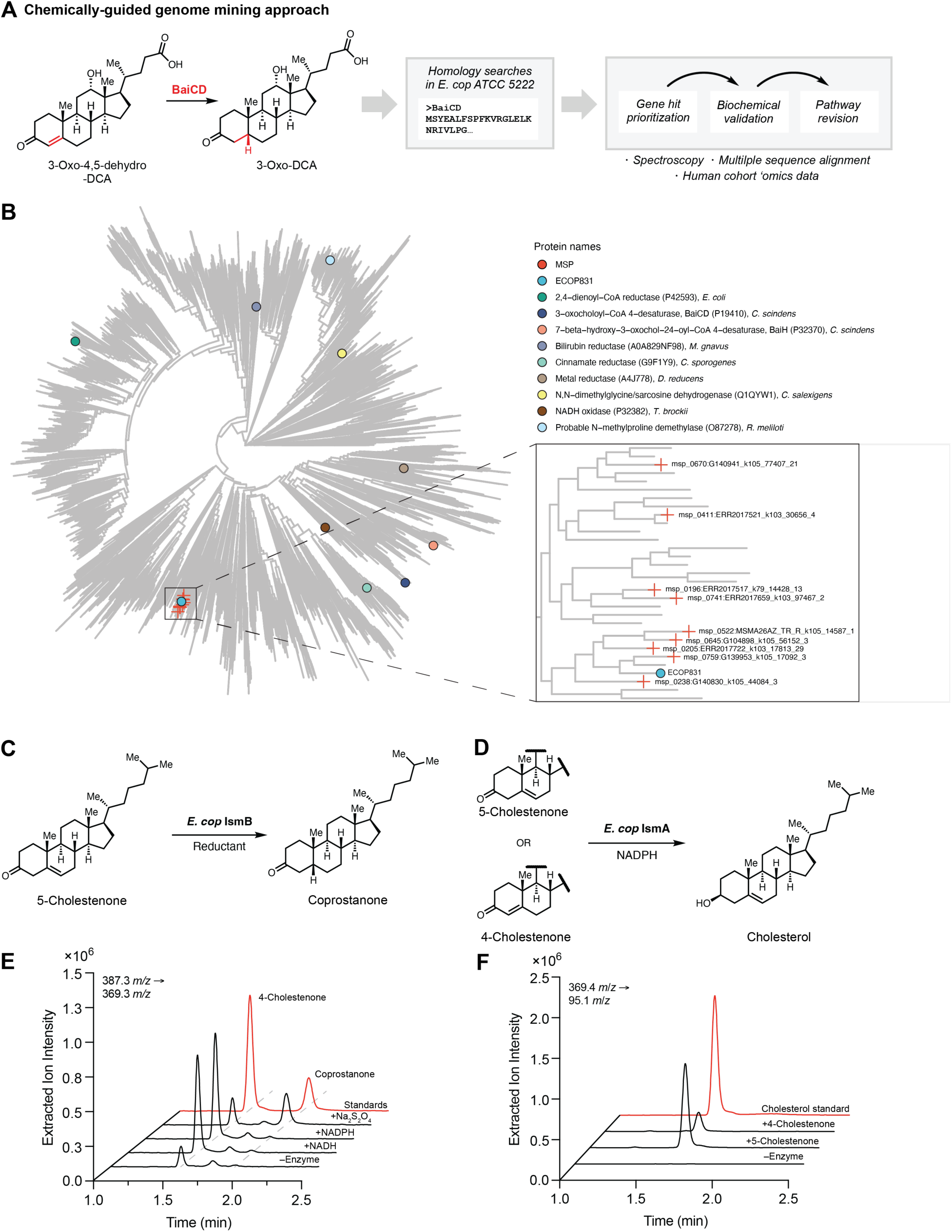
Chemically guided genome mining approach led to the discovery of 5-cholestenone reductase IsmB. **(A)** BaiCD from *C. scindens* was used to guide genome mining in *E. cop* ATCC 51222 for the missing gene(s) and enzyme(s) in gut microbial coprostanol production. **(B)** Fe–S flavoenzyme superfamily phylogenetic tree was constructed using FastTree^40^ from 9,440 non-redundant protein sequences across the bacterial kingdom, and includes 9 clades containing previously experimentally characterized proteins and/or enzymes listed in UniProt that are indicated in circles. All sequences used as inputs contained Pfam domains Oxidored_FMN (PF00724) and Pur_redox_2 (PF07992) and were retrieved from UniprotKB (Accessed July 07, 2025). **(C)** Chemical reaction catalyzed by *E. cop* IsmB using 5-cholestenone as the substrate and different chemical reductants. **(D)** LC–MS/MS extracted ion chromatograms (EICs) of the 369.3 *m/z* fragment ion of 16 h anaerobic incubations of 5-cholestenone with purified recombinant *E. cop* IsmB and different reductants reveal that 5-cholestenone is efficiently converted to coprostanone only when sodium dithionite (Na_2_S_2_O_4_) is used as the reductant. Any non-reacted 5-cholestenone substrate isomerizes to 4-cholestenone and accumulates over time. **(E)** Chemical reaction scheme of the first step of the *ism* pathway in the reverse direction catalyzed by *E. cop* IsmA, the conversion of 4- or 5-cholestenone to cholesterol. **(F)** LC–MS/MS EICs of the 95.1 *m/z* fragment ion of cholesterol produced in 16 h anaerobic incubations of 4- or 5-cholestenone with purified recombinant *E. cop* IsmA. These assays reveal that 5-cholestenone is preferentially converted to cholesterol over 4-cholestenone, when NADPH is used as the cofactor, and indicates that 5-cholestenone is an on-pathway intermediate. (C–F) n = 3 biologically independent replicates.

Because the substrate of 3-oxo-4,5-dehydro DCA structurally resembles 4-cholestenone, we hypothesized that the *ism* pathway might involve an Fe–S cluster-dependent flavoenzyme homologous to BaiCD that catalyzes the stereoselective two electron reduction of 4-cholestenone to coprostanone. We searched the genome of *E. cop* using *C. scindens* BaiCD as a query, and identified a potential candidate, ECOP831 (29% amino acid (AA) identity (ID), 100% query coverage, and 44% similarity to *C. scindens* BaiCD) (**Figure 2A**). Analysis of the ECOP831 protein sequence with InterProScan^31^ revealed that this protein belongs to the NADH:flavin oxidoreductase (IPR051793) and the 2,4-dienoyl-CoA reductase (2,4-DCR) (PTHR42917) families and identified two distinct domains: an N-terminal flavin mononucleotide (FMN)-binding old yellow enzyme (OYE)/oxidoreductase domain (IPR001155) and a C-terminal flavin adenine dinucleotide (FAD)/nucleotide adenine dinucleotide (NAD)-binding domain (IPR023753). Notably, the InterProScan analysis did not reveal domains belonging to Fe–S cluster-binding protein families. Since *C. scindens* BaiCD was reported to be an [4Fe–4S]-binding flavoenzyme and is homologous to ECOP831^30^, we suspected that ECOP831 may also contain an Fe–S cluster binding domain. Additionally, when the ECOP831 gene was mapped onto the genome of *E. cop*, we found that it is not colocalized with *ismA* in *E. cop*’s genome. We found *ismA* is not flanked by any regulatory genes or genes annotated with functions that may be related to sterol metabolism (**Figure S1B**).

We also used *C. scindens* BaiN as a search query in our chemically-guided genome mining approach, as BaiN is reported to catalyze the reduction of 3-oxo-4,5-dehydro DCA to 3-oxo-DCA.^32^ The BaiN enzyme family remains poorly characterized, but InterProScan revealed it to contain an FAD binding domain (IPR036188), agreeing with the yellow color of the recombinantly expressed protein.^32^ We found that *E. cop* encodes two homologs of BaiN, ECOP530 and ECOP884, and considered them as potential candidates for the missing enzyme in the conversion of cholestenone to coprostanone by *E. cop*.

To assess if ECOP831 is a member of the greater bacterial Fe–S flavoenzyme superfamily, we reconstructed a Fe–S flavoenzyme maximum-likelihood phylogenetic tree that was first reported by Andreu and coworkers (**Figure 2B**).^33^ We identified and included 9,440 protein sequences that contain PFAMs PF00724 and PF07992 and are between 600–800 AA in length. Within the phylogenetic tree, we annotated 10 clades based on experimentally characterized proteins and/or enzymes listed in the UniProt database. We found a distinct clade that contains ECOP831 and proteins homologous to ECOP831 from human stool MSPs (**Figure 2B**), with the vast majority of the clades in the phylogenetic tree containing proteins with unknown functions.

We next predicted the structure of ECOP831 using AlphaFold3^34^ with high confidence scores and low predicted alignment errors (**Figure S2A**) and aligned it with the X-ray protein crystal structure of *E. coli* 2,4-DCR,^35^ which contains an N-terminal-bound FMN cofactor, a C-terminal-bound FAD cofactor, a fully occupied [4Fe–4S] cluster, and the cognate 2,4-dienoyl-CoA thioester substrate bound in the N-terminal OYE domain (**Figure 3A** and **S2B**). This structure alignment was obtained with a low root mean squared deviation (RMSD) of 1.207 Å and revealed both flavin cofactor-binding domains and a putative centrally located ferredoxin-like Fe–S cluster binding domain consisting of Cys386, Cys388, Cys392 and Cys408 in ECOP831. This structure alignment also led us to hypothesize that ECOP831 may bind 4-cholestenone in its N-terminal OYE/oxidoreductase domain. We then obtained the AlphaFold3 predicted structure of *C. scindens* BaiCD and aligned it with the ECOP831 AlphaFold structure (RMSD 1.708 Å) (**Figure 3B**). This highlighted the predicted centrally-located, conserved putative [4Fe–4S] cluster binding domain in both enzymes (**Figure S2C**). Altogether, these analyses revealed that *E. cop* encodes a candidate enzyme homologous to BaiCD that potentially binds flavin cofactors and an Fe–S cluster.

**Figure 3.**
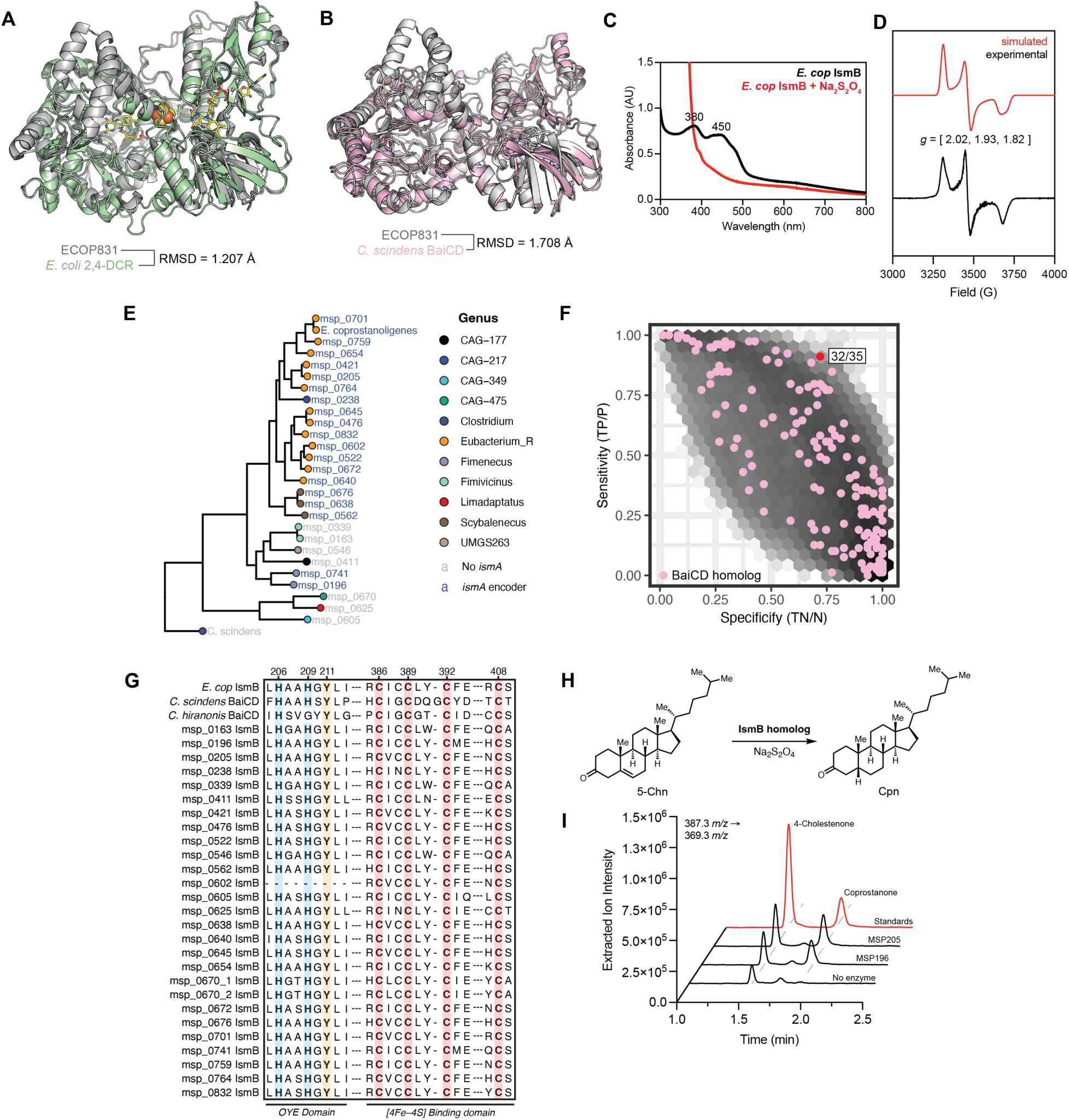
IsmB is an Fe–S cluster- and flavin-dependent metalloenzyme encoded in uncultured human gut bacteria. AlphaFold3 predicted structure of ECOP831 (gray) aligned to **(A)** the X-ray crystal structure of *E. coli* 2,4-DCR (PDB 1PS9) and **(B)** to the AlphaFold3 predicted structure of *C. scindens* BaiCD. **(C)** UV–vis spectroscopy suggests that IsmB binds flavin cofactor(s) with absorption maxima at 380 and 450 nm. **(D)** EPR spectroscopy at 10 K of *E. cop* IsmB reveals a rhombic spectrum and indicates the presence of a protein bound reduced [4Fe–4S] cluster. **(E)** 19 different MSPs containing both *ismB* and *ismA* genes were identified in human gut microbiome datasets. Phylogenetic tree was created using GTDB-tk^58^ and includes all IsmB-encoding MSPs as well as *E. cop*. **(F)** Scores of specificity and sensitivity in relation to presence of coprostanol in stool metabolomes were calculated for each cluster of homologous proteins present in stool metagenomes from the PRISM and HMP2 datasets, which were queried with the biochemically characterized *C. scindens* BaiCD that performs a similar chemical transformation. 32 of 35 IsmB homologs identified in the BLAST search fell into the same protein cluster with high sensitivity and specificity for stool coprostanol, while the remaining three fell into protein clusters displaying high specificity but low sensitivity for coprostanol. **(G)** Multiple sequence alignment of IsmB homologs from *E. cop* and human gut metagenomic species with BaiCD homologs from *C. scindens* and *C. hiranonis* highlights highly conserved N-terminal His and Tyr amino acid residues and a Cys-X_2_-Cys-X_2_-Cys-X_15_-Cys 4Fe–4S cluster binding motif. **(H)** Chemical reaction catalyzed by IsmB homologs from human gut MSPs. **(I)** LC–MS/MS EIC of the 369.3 *m/z* fragment ion of coprostanone produced in 16 h anaerobic incubations of 5-cholestenone with purified recombinant IsmB homologs from by MSPs 196 and 205. (G–H) n = 3 biologically independent replicates.

### Biochemical characterization of the putative sterol-metabolizing microbial Fe–S flavoenzyme

To determine if the putative Fe–S flavoenzyme ECOP831 can reduce 4-cholestenone to coprostanone, we expressed it in *E. coli* co-expressing the *E. coli iscSUA-HscBA-Fd* operon on a pPH149 plasmid^36^ and purified the enzyme using Ni^+^-affinity chromatography under strict anaerobic conditions. We observed a ∼75.5 kDa molecular weight band in the imidazole elution fractions after SDS-PAGE analysis (**Figure S3A**). The purified enzyme was yellow-brown colored, and size exclusion chromatography (SEC) analysis revealed that ECOP831 exists as a monomer in solution (**Figure S3B**). Next, we tested the activity of ECOP831 toward the conversion of 4-cholestenone to coprostanone under anaerobic conditions using NADH, NADPH, or sodium dithionite (Na_2_S_2_O_4_) as the candidate reductants. Using LC–MS/MS quantitation, only trace activity of ECOP831 toward 4-cholestenone was observed, even when stoichiometric ratios of purified ECOP831 were used in the assays and when compared to a no enzyme control (**Figure S3C**). This finding was surprising and unexpected considering our previous observation that *E. cop* IsmA produced 4-cholestenone as the only detected product^12^, and the fact that all previously biochemically characterized steroid-and bile acid-metabolizing Fe–S flavoenzymes accept α,β-unsaturated cognate substrates.^30,37,38^ The more recently discovered class of gut microbial Fe–S cluster and flavin dependent 5β-reductases, Ci2350 and OsrB, exhibited broad substrate promiscuity towards several structurally distinct α,β-unsaturated steroid substrates, but displayed trace or no detectable activity toward 4-cholestenone.^37,38^ These reports along with our initial finding point to a possible difference in substrate preference for the gut microbial enzymatic reduction of cholestenone to coprostanone that we hypothesize to proceed through an unsaturated 4-keto intermediate.

We then tested if ECOP831 could reduce the unsaturated Δ^5,6^ bond in cholesterol to generate coprostanol directly, but no activity was detected using LC–MS/MS (**Figure S3D**). These findings further support the proposal that IsmA must first oxidize cholesterol to produce a 3-keto intermediate. We hypothesized that, although 4-cholestenone was proposed to be the on-pathway intermediate for the gut microbial conversion of cholesterol to coprostanol, it may not be the preferred substrate isomer for ECOP831.

We next investigated whether the BaiN homologs, ECOP530 and ECOP884, catalyze the conversion of 4-cholestenone to coprostanone. We aerobically expressed each candidate protein in *E. coli*, and aerobically purified them using Ni-NTA affinity chromatography, which produced yellow colored proteins, indicating both proteins bind flavin cofactors. Upon assaying the enzymes under aerobic conditions, and using either NADH, NADPH or Na_2_S_2_O_4_ as reductants, we observed that neither ECOP530 nor ECOP884 catalyzed the conversion of 4-cholestenone to coprostanone (**Figure S3E**). This result further supports our hypothesis that the second step in the microbial conversion of cholesterol to coprostanol may not proceed through a 4-cholestenone intermediate.

5-cholestenone is the immediate product of the enzymatic oxidation of cholesterol by IsmA and has also been proposed as an intermediate in coprostanol formation.^22^ Thus, we considered the possibility that ECOP831 may convert 5-cholestenone to coprostanone. When ECOP831 was incubated with 5-cholestenone in the presence of Na_2_S_2_O_4_ under anaerobic conditions, high levels of conversion to coprostanone were observed (**Figure 2C** and **2D**). Additionally, in the assays that omitted the enzyme and/or Na_2_S_2_O_4_, 4-cholestenone was detected as the major product, indicating that 5-cholestenone readily isomerizes to 4-cholestenone under the assay conditions (**Figure 2D**). The rate of isomerization was found to be 5-fold faster in assays that included ECOP831 without reductant, indicating that the enzyme may catalyze the isomerization of 5-cholestenone to 4-cholestenone. This isomerization reaction had previously been reported when 5-cholestenone was added to liquid cultures of coprostanol-producing stool samples as a potential substrate^39^ and was reported to be accelerated in the presence of a Brønsted acid or base.^40^ In our assay conditions, we hypothesize that the 5β-reductase activity of ECOP831 is kinetically faster than rate of isomerization, and thus we detect higher levels of coprostanone rather than the isomerization product 4-cholestenone. Our observation that ECOP831 selectively reduces 5-cholestenone but not 4-cholestenone into coprostanone, and that 5-cholestenone isomerizes to 4-cholestenone in the absence of enzyme implies that the detection of 4-cholestenone in previous work studying the gut microbial cholesterol-to-coprostanol conversion and in our initial characterization of IsmA may have been a result of non-specific acid–base-catalyzed isomerization.

Lastly, we tested if the BaiN homologs, ECOP530 and ECOP884, may also convert 5-cholestenone to coprostanone. Using clarified lysates from *E. coli* heterologously expressing either protein in LC–MS-based assays with and without additional FAD cofactor and using either NADH, NADPH or Na_2_S_2_O_4_ as reductants, we observed no conversion of 5-cholestenone to coprostanone by ECOP530 or ECOP884 (**Figure S3F**). This result indicated that the enzymatic reduction of 5-cholestenone to coprostanone is specific to ECOP831. With the activity of ECOP831 toward the reduction of 5-cholestenone to coprostanone biochemically confirmed, we named it IsmB.

### Both 4- and 5-cholestenone intermediates are relevant in the *ism* pathway

The relevancy of 5-cholestenone as an on-pathway intermediate and substrate for the 5β-reductase responsible for producing coprostanone has been largely unexplored. To investigate if IsmB works productively with IsmA in the *ism* pathway and whether 4- or 5-cholestenone are implicated in coprostanol production, we incubated IsmA with each potential intermediate and measured the ability of IsmA to catalyze the conversion of 4- and 5-cholestenone to cholesterol, i.e., the first step of the *ism* pathway in the reverse direction. We observed that IsmA efficiently catalyzes the stereoselective reduction of 5-cholestenone to 3β-configured cholesterol (**Figures 2E and 2F**). IsmA also isomerized and reduced 4-cholestenone to cholesterol, albeit with 6-fold lower activity compared to assays using 5-cholestenone (**Figure 2F**). Collectively, these results imply that out of the two possible isomer intermediates that are hypothesized to be on-pathway in coprostanol production, 5-cholestenone is the preferred substrate for IsmB.

To further investigate IsmB’s preference for 5-cholestenone over 4-cholestenone, we performed molecular docking analyses using the *E. cop* IsmB AlphaFold3-predicted structure and the diffusion generative model DiffDock.^41^ Either 4-cholestenone or 5-cholestenone was docked into the *E. cop* IsmB predicted structure, and the docking models were inspected for potentially important protein amino acid–substrate interactions (**Figures S4A and S4B**). We found that both 4-and 5-cholestenone are predicted to bind *E. cop* IsmB in the N-terminal FMN-binding OYE-like domain and within proximity of His206, His209 and Tyr211, supporting our hypothesis that it serves as the enzyme active site. However, we could not determine from these molecular docking models whether *E. cop* IsmB preferentially binds either 4- or 5-cholestenone.

To further assess the relevance of IsmB’s observed specificity *in vitro* to coprostanol formation in bacterial cultures, we prepared *E. cop* liquid cultures incubated with either cholesterol, 4-cholestenone, 5-cholestenone, or coprostanone (**Figure S4C**). The *E. cop* culture incubated with cholesterol accumulated trace amounts of 4-cholestenone and coprostanone in the span of 96 hours, demonstrating these intermediates are short-lived. Additionally, coprostanone was converted to coprostanol in the cultures fed coprostanone. Unlike the IsmB enzyme assays, we observed that *E. cop* converted both 4- and 5-cholestenone to coprostanol over the span of 96 hours, in agreement with previous work.^22^ This result implies two scenarios occurring in *E. cop* bacterial cultures: (i) *E. cop* encodes an unidentified 5β-reductase that has substrate specificity for 4-cholestenone or (ii) *E. cop* converts 4-cholestenone back to 5-cholestenone using an unidentified isomerase domain or protein. Our observation that IsmA catalyzes the isomerization and reduction of 4-cholestenone back to cholesterol preliminarily supports the latter scenario. Collectively, these results revealed that 5-cholestenone is the preferred substrate of IsmB in purified enzyme assays, and that 5-cholestenone is a relevant intermediate in the gut microbial conversion of cholesterol to coprostanol, thus preliminarily revising the *ism* pathway.

### IsmB binds redox-active flavin and Fe–S cluster cofactors

The substrate specificity of IsmB and our observation through bioinformatic analysis that the vast majority of the members of bacterial Fe–S flavoenzyme superfamily are uncharacterized prompted us to further investigate the cofactors and cofactor-binding domains of this enzyme. Since the enzymes BaiCD and 2,4-DCR bind FMN in their N-terminal domain and FAD in their C-terminal domain, we hypothesized that IsmB also binds FMN and FAD cofactors in both its N- and C-terminal domains, respectively. To test this, we first collected the UV–vis spectrum of IsmB under anaerobic conditions and observed two absorbance features at 380 and 450 nm (**Figure 3C**), which are commonly found in the UV–vis spectra of many flavin dependent proteins^42^ and specifically in enzymes containing OYE-like domains that are homologous to *E. cop* IsmB.^43–45^ When IsmB was incubated with excess Na_2_S_2_O_4_, we observed the disappearance of the absorbance maxima at 380 and 450 nm, suggesting the enzyme was isolated with its flavin cofactors in their oxidized states, and that the reduction of the oxidized FMN or FAD in IsmB by Na_2_S_2_O_4_ converts the flavin cofactors into their reduced semiquinone state.^46,47^ We then assessed the identity of the flavin cofactors that IsmB binds using a flavin extraction HPLC–diode array-based and quadrupole time-of-flight (Q-TOF) high resolution mass spectrometry (HRMS)-based assays. We found that IsmB binds both FMN and FAD in an approximate 1:1:1 IsmB monomer-to-FMN-to-FAD ratio (**Figure S5A–C**), and we detected [M−H]^−^ ion counts for both FMN (455.097 *m*/*z*) and FAD (784.150 *m*/*z*) for IsmB relative to analytical standards (**Figure S5D–E**). This finding further supports that IsmB contains FMN- and FAD-binding domains and is similar to other enzymes in this family.

To determine which domains likely bind the FMN or FAD cofactors in IsmB, and because *E. coli* 2,4-DCR is the only member of Fe–S flavoenzyme superfamily whose protein crystal structure has been solved, we turned to the multi-modal foundation model for protein structure prediction, Chai-1.^48^ We obtained the predicted protein structure of *E. cop* IsmB bound to FMN and FAD ligands, and found that *E. cop* IsmB is predicted to bind FMN in the N-terminal OYE/oxidoreductase domain and FAD in the C-terminal FAD/NAD-binding domain (**Figure S5F)**. These insights suggest that IsmB uses both flavin cofactors as part of its electron transfer mechanism to generate a fully reduced FMN cofactor in the N-terminal active site, which may then reduce 5-cholestenone through hydride delivery. Similar electron transfer mechanisms involving FAD, 4Fe–4S, and FMN have been proposed for the structurally-related *E. coli* 2,4-DCR^35^ and the N47 enrichment culture-encoded 2-napththoyl-CoA reductase (NCR), in which a centrally located 4Fe–4S cluster helps mediate electron transfer between FAD and FMN.^46^

We then focused on characterizing the putative Fe–S cluster predicted to bind in the central ferrodoxin-like domain of *E. cop* IsmB. We first quantified the molar ratios of iron (Fe^2/3+^) and sulfide (S^2−^) in IsmB using inductively-coupled plasma–mass spectrometry (ICP–MS) and a modified sulfide colorimetric assay. We observed that *E. cop* IsmB contained 5.08 ± 0.32 moles of Fe^2/3+^ and 4.03 ± 0.07 moles of S^2−^ per enzyme monomer, suggesting the presence of at least one [4Fe–4S]^2/3+^ cluster (**Figure S5G**). Electron paramagnetic resonance (EPR) spectroscopy was employed to further characterize the Fe–S cluster in *E. cop* IsmB. Using excess Na_2_S_2_O_4_ as the reductant, the *E. cop* IsmB produced a rhombic EPR spectrum with *g* values of [2.02, 1.93, 1.82], indicating that *E. cop* IsmB contains a redox active [4Fe–4S]^2/3+^ cluster (**Figure 3D**). In all, these analyses demonstrate that *E. cop* IsmB is a redox active flavin- and Fe–S cluster-dependent metalloenzyme whose cofactors may play important roles in the reduction of 5-cholestenone to coprostanone.

Our findings that *E. cop* IsmB does not efficiently reduce 5-cholestenone to coprostanone in the presence of the nicotinamide-based reductants NADH and NADPH, and that enzyme activity is observed only with Na_2_S_2_O_4_ as the reductant provide evidence that *E. cop* IsmB requires a strongly reducing low-redox potential electron donor. In the case of NCR, a ferredoxin or a ferredoxin-like domain of an unknown oxidoreductase (*E’* ≈ −500 mV) is proposed to be the electron donor in its native organism N47.^43,46^ Recently, the human gut bacterial Fe–S cluster and flavin-dependent 5β-reductase, *C. steroidoreducens* OsrB, that metabolizes natural and synthetic steroid hormones was found to require the strong abiological reductant and electron mediator methyl viologen (*E’* ≈ −446 mV^51^) in order to detect activity.^38,52^ It is therefore likely that *E. cop* IsmB requires a protein-based electron donor to reduce its FMN cofactor and carry out the reduction of 5-cholestenone to coprostanone under native conditions. The FAD binding domain of the *E. cop* IsmB AlphaFold3 predicted structure contains a solvent-exposed cleft, which may serve as the binding site of a putative redox partner.

### Uncultured human gut bacteria associated with coprostanol production encode IsmB homologs

Our previous finding that IsmA-encoding uncultured human gut bacteria are detected in human stool metagenomes and strongly correlate with coprostanol presence in stool metabolomes^12^ prompted us to investigate if homologs of *E. cop* IsmB are encoded by these uncultured human gut bacteria. To validate the relevance of IsmB in gut microbial cholesterol metabolism in humans, *E. cop* IsmB was used to query gut bacterial MSPs from several human participant data sets, including the Framingham Heart Study (FHS), the Prospective Registry in IBD Study at MGH (PRISM)^53^, Integrative Human Microbiome Project (HMP2)^54^, CVON^55^, 500 Functional Genomics (500FG) cohort^56^, and Jie et al.^57^ Using a 50% AA ID cutoff, we found 35 protein sequences homologous to *E. cop* IsmB (**Figure S6**). Of the 35 IsmB homologs, 19 were from MSPs that also encode an IsmA homolog with at least 50% AA ID to *E. cop* IsmA (**Figure S6**). The majority of these MSPs are of the genus Eubacterium, with other genera represented being Fimenecus and Scybalenecus (**Figure S6** and **Figure 3C**). In an analysis of paired human gut metagenomes and stool metabolomic samples, we found one protein family cluster was highly discriminative for the presence of coprostanol (**Figure 3F**), consistent with a role in coprostanol production. 32 of the 35 IsmB homologs identified fell into this protein cluster, with the other three homologs falling into other protein clusters that lacked sensitivity for coprostanol in associated stool metabolomes.

To aid in assessing the bacterial taxonomy of the MSPs encoding IsmB homologs, we analyzed their phylogenetic relationship to known microbial isolates using a set of single-copy marker genes. We found the IsmA/IsmB-encoding MSPs and *E. cop* form a distinct subclade separate from the MSPs that encode IsmB homologs but not IsmA homologs (**Figure 3E**). Two exceptions to this subclade separation are MSP 196 and MSP 741, which each encode IsmA and IsmB homologs, but do not clade together with the other *ismA/B*^+^ MSPs. This observation suggests that MSP 196 and MSP 741 may have evolved separately from the clade of uncultured bacteria that predominantly encode both *ism* genes. Unsurprisingly, the IsmB homologs encoded by MSP 196 and MSP 741 share the lowest % nucleotide (NT) ID with *E. cop* IsmB, having 59% and 58% NT ID, respectively. Upon generating a protein phylogenetic tree of IsmB homologs, we once again observed the clustering of IsmB homologs from IsmA-encoding MSPs into a distinct clade, and the separation of the IsmB homologs encoded by MSPs 196 and 741 (**Figure S7A**). We next performed cophylogenetic (cophylo) analysis of the 19 IsmA and IsmB homologs and observed clear coevolutionary relationships (**Figure S7B**). The low degree of entanglement observed in the cophylo tree suggests a low degree of incongruence between the evolution of IsmA and IsmB in coprostanol-producing bacteria.

To investigate if the IsmB homologs from the human gut MSPs contained the predicted catalytic and cofactor binding domains, we aligned the protein sequences of IsmB homologs from *E. cop* and human gut MSPs along with BaiCD homologs from *C. scindens* and *C. hiranonis* (**Figure 3G**). This analysis revealed high homology around the N-terminal FMN-binding OYE-like domains, including two conserved His residues, H206 and H209, and a putative catalytic Tyr, Y211, in *E. cop* IsmB. These conserved residues have been found to be important for substrate binding and catalysis in OYEs and in enzymes containing OYE-like domains.^34,42,44,58^ Additionally, all IsmB homologs contain the centrally-located Cys-X_2_-Cys-X_2_-Cys-X_15_-Cys Fe–S cluster binding motif, further indicating that this domain and metallocofactor is an essential electron-transfer mediator in this broader enzyme family. To assess the difference in the protein folds, two IsmB homologs with varying differences in AA % ID relative to *E. cop* IsmB were further investigated (ERR2017517_k79_14428_13 from MSP 196 and ERR2017722_k103_17813_29 from MSP 205 with 59% and 70% AA ID, respectively, to *E. cop* IsmB). When the AlphaFold3 predicted structure of MSP 196 IsmB was aligned onto the *E. cop* IsmB predicted structure, the RMSD was found to be 0.511 Å, indicating that their predicted protein folds are highly similar and conserved (**Figure S8A**). Similarly, when the AlphaFold3 predicted structures of MSP 205 IsmB and *E. cop* IsmB were aligned, an even lower RMSD of 0.326 Å was obtained, and thus agreeing with the greater % AA ID shared between the two proteins (**Figure S8C**). These insights demonstrate that although the IsmB homologs from uncultured bacteria from different phylogenetic subclades have varying levels of shared AA ID, their predicted protein folds are highly conserved, suggesting that their biochemical functions may also be conserved.

To investigate the ability of the human gut bacterial IsmB homologs to convert 5-cholestenone to coprostanone, and thus validate their relevance to coprostanol production by human gut bacteria, we turned to biochemical characterization. The IsmB homologs from MSP 196 and MSP 205 were first recombinantly overexpressed in *E. coli* and purified under strict anaerobic conditions. Size exclusion chromatography revealed that, similar to *E. cop* IsmB, the human gut bacterial homologs also exist as monomers in solution (**Figure S8E** and **S8F**). Further characterization using Q-TOF HRMS (**Figures S5D and S5E**), UV–vis spectroscopy and EPR (**Figure S8G** and **S8H**) revealed mass spectrometry and spectroscopic features indicating that the MSP 196 and MSP 205 IsmB homologs also bind redox-active FMN, FAD and Fe–S cluster cofactors. Upon assaying the enzymes using LC–MS, we observed that both IsmB homologs from MSP 196 and MSP 205 efficiently converted 5-cholestenone to coprostanone when incubated anaerobically with catalytic amounts of enzyme and Na_2_S_2_O_4_ as the reductant (**Figures 3H** and **3I**). When the IsmB homologs from MSP 196 and MSP 205 were assayed using 4-cholestenone, we observed no conversion to coprostanone, and revealing that these enzymes possess the same specificity for 5-cholestenone as *E. cop* IsmB (**Figure S8I and S8J**). This is the first time, to our knowledge, that human gut microbial enzymes responsible for the second step in the production of coprostanol have been identified and biochemically characterized.

### Site-directed mutagenesis of *E. cop* IsmB reveals the importance of the conserved amino acid residues

Observing that human gut bacterial homologs of IsmB contain the highly conserved N-terminal His and Tyr residues postulated to be involved in substrate binding and catalysis, and the putative Fe–S cluster tetra Cys binding motif, we used site-directed mutagenesis to further probe their importance for converting 5-cholestenone to coprostanone. After protein expression in *E. coli* and anaerobic purification, the activity of each variant was tested using endpoint enzyme assays. When the His and Tyr residues in the N-terminal OYE-like domain of IsmB were mutated, the IsmB variants displayed moderately lower enzyme activity when compared to the WT *E. cop* IsmB in 16 hour endpoint assays (**Figure S9B** and **S9C**). Specifically, the H206A and H209A single and H206A/H209A double variants produced 216 µM, 213 µM, and 210 µM coprostanone, respectively, compared to 237 µM coprostanone for the WT enzyme, suggesting a modest drop in catalysis. In the case of the Y211F variant, a larger drop in activity was observed, with 162 µM coprostanone being produced. Unlike the Y211F variant, the Y211A variant did not lead to a large drop of coprostanone production. While these results were initially surprising, the modest changes in the enzyme activity of these IsmB active site variants recapitulate the activities seen with *Saccharomyces carlsbergensis* OYE1 active site variants, where mutating the active site Tyr196 to Phe and His191 to Asn did not result in a loss of catalysis and instead led to a change in the enzyme kinetic properties.^58,59^ Similarly, when the predicted catalytic Tyr167 of the progesterone-metabolizing 5β-reductase Ci2350 was mutated to Ala, a drop in the production of 5β-dihydroprogesterone was observed instead of a complete loss of activity.^37^ The hydrophobicity and poorly solubility of 5-cholestenone in aqueous media precluded completing the enzyme kinetic characterization of IsmB, and thus we cannot determine how mutations to IsmB’s active site AA residues affect its enzyme kinetic properties.

We next tested the single point *E. cop* IsmB mutants targeting the tetra cysteine binding motif, C386A, C389A, C392A and C408A, using endpoint assays. We unexpectedly did not observe a complete loss of enzyme activity. The IsmB variants C386A, C389A, C392A and C408A produced 177 µM, 161 µM, 173 µM, and 209 µM coprostanone, respectively, compared to 237 µM coprostanone for WT IsmB (**Figure S9B** and **S9C**). This observation may result from the direct reduction of FMN in the OYE domain of IsmB by Na_2_S_2_O_4_, and thus bypassing the electron transfer pathway that may involve the 4Fe–4S cofactor.^35,43,46^ Altogether, these results reveal that although these amino acid residues are highly conserved, they are not essential for IsmB enzyme activity under the in vitro assay conditions. The highly conserved amino acid residues in the IsmB active sites may play essential roles in enzyme catalysis using the native reductant or electron donor and/or under in vivo conditions.

To examine how substitution of these conserved amino acid residues may affect cofactor binding, we characterized the IsmB variants using UV–vis spectroscopy and quantified the occupancy of their Fe–S clusters. Perturbations to the residues Y211, H206 and H209 in the OYE-like domain resulted in little changes to the UV–vis spectral feature compared to the WT *E. cop* IsmB (**Figure S9D**). Unexpectedly, the IsmB variants containing mutations to the conserved Fe–S cluster binding domain produced UV–vis spectra with altered features compared to the absorption spectrum of WT *E. cop* IsmB (**Figure S9E**). Specifically, the *E. cop* IsmB variant C386A produced a UV–vis spectrum with diminished absorbance maxima at 380 and 450 nm, indicating that this perturbation results in reduced flavin cofactor binding (**Figure S9E**). We next quantified both the Fe–S cluster in the *E. cop* IsmB variants by performing ICP–MS and colorimetric sulfide-content assays to quantify the moles of Fe^2/3+^ and S^2−^. We observed a drop in Fe^2/3+^ and S^2−^loading for the H206 and H209 IsmB variants, with H209 containing nearly half of the Fe^+2/3^ as compared to WT *E. cop* IsmB (**Figure S9F**). The Fe–S cluster assembly in the Y211A and Y211F variants were weakly impacted, as observed by their molar ratios of Fe and S being similar to WT *E. cop* IsmB. Unsurprisingly, we observed that the mutations to the Fe–S cluster binding domain led to significant drop in Fe^2/3+^ and S^2−^ loading in those *E. cop* IsmB variants. The variant C386A was the most impacted, containing 0.84 ± 0.11 Fe^2/3+^ and 0.73 ± 0.02 S^2−^, indicating that this amino acid residue is sensitive to any structural perturbations and greatly affects the assembly and chelation of the [4Fe–4S] cluster (**Figure S9F**). These results demonstrate how perturbations to the amino acid sequence in the IsmB active site and cofactor binding sites results in changes proximal and distal to those sites in regard to cofactor loading and binding.

### Cholesterol induces the expression of *ismA* while *ismB* is constitutively expressed in *E. cop*

In an effort to understand the expression of *ismA* and *ismB* in *E. cop* and how the transcription of these genes translates to enzymatic activity, we assayed *E. cop* lysates prepared from cultures that were grown in the presence or absence of cholesterol. To first test if the expression of *ismA* is induced by cholesterol, *E. cop* lysates were assayed for IsmA activity using cholesterol as a substrate and in the presence of NADP^+^ as the cofactor. The production of 4-cholestenone was detected only with lysates prepared from *E. cop* cultures grown in presence of cholesterol (**Figure 4A**), and there was also no conversion of cholesterol to 4-cholestenone detected in the assays using lysates from *E. cop* cultures grown in the absence of cholesterol and treated with a vehicle control (**Figure 4B**). In this case, 5-cholestenone could not be detected as we hypothesize that the assay conditions isomerizes the 5-cholestenone produced by IsmA to 4-cholestenone before the LC–MS/MS quantitation is completed. These results indicate that the expression of *ismA*, and thus the enzymatic activity of IsmA, is induced by cholesterol, in agreement with our previous observation.^12^

**Figure 4.**
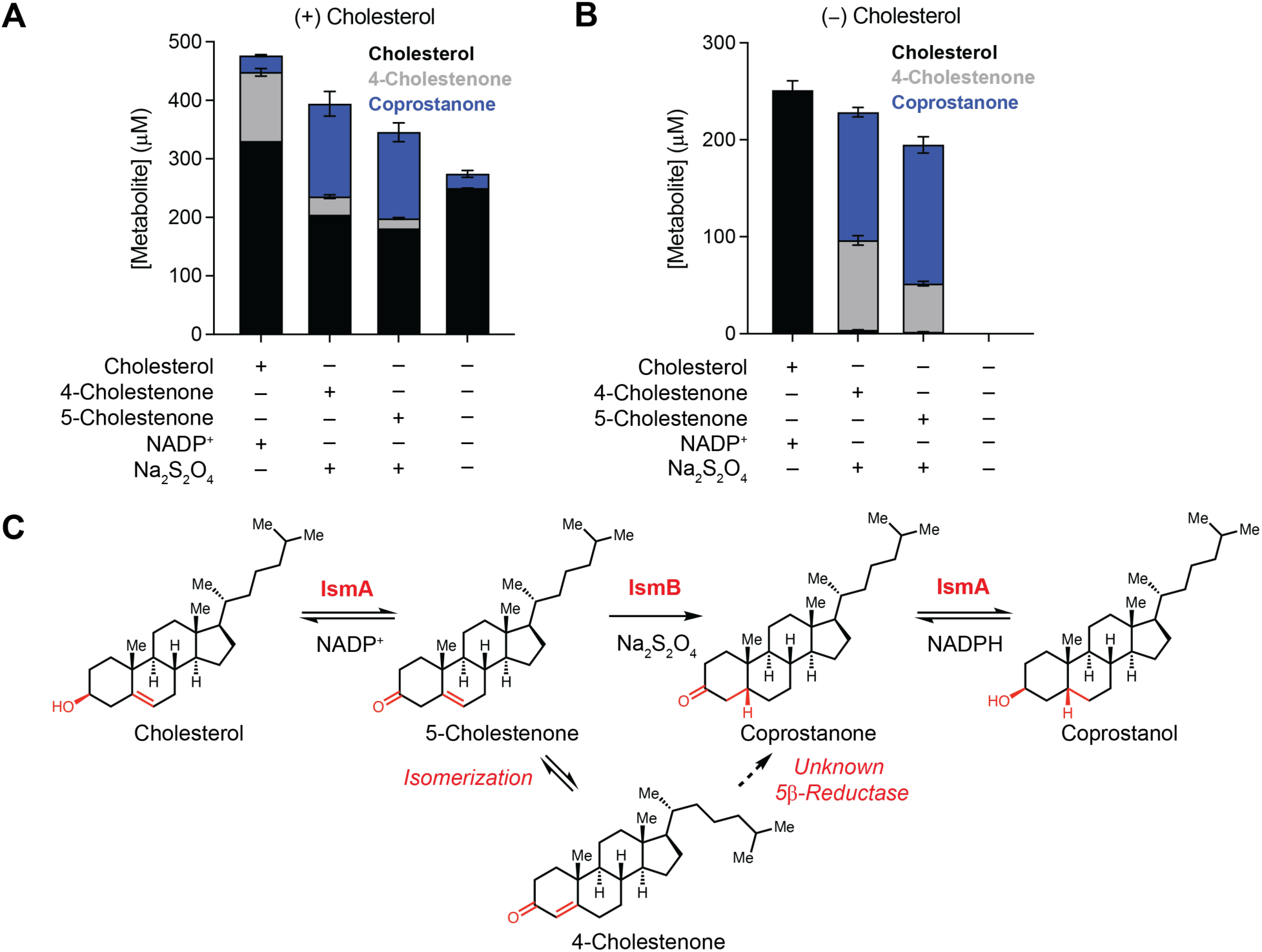
Cholesterol has different impacts on the expression of IsmA and IsmB in *E. coprostanoligenes* ATCC 51222. **(A)** Assays using lysates from *E. coprostanoligenes* ATCC 51222 cultures supplemented with 500 µM cholesterol at an OD_600_ = 0.5. Error bars represent the standard error mean (SEM) of three biological replicates. **(B)** Assays using lysates from *E. coprostanoligenes* ATCC 51222 cultures treated with vehicle at an OD_600_ = 0.5. Error bars represent the standard error mean (SEM) of three biological replicates. **(C)** Revised *ism* pathway for the gut microbial conversion of intestinal cholesterol to coprostanol that involves the reduction of the intermediate 5-cholestenone by IsmB to generate the intermediate coprostanone. (A–B) n = 3 biologically independent replicates, data presented are the ± standard error mean (SEM).

To test whether the expression of *ismB* is induced by cholesterol, we performed *E. cop* lysate assays with 4- and 5-cholestenone as substrates along with Na_2_S_2_O_4_ as the reductant. We observed the conversion of both 4- and 5-cholestenone to coprostanone in assays using lysates from both *E. cop* cultures grown with and without cholesterol (**Figures 4B and 4C**).The unexpected activity for both 4- and 5-cholestenone observed in these *E. cop* lysates did not recapitulate the substrate specificity for 5-cholestenone observed in assays with purified *E. cop* IsmB, but did agree with prior results from *E. cop* cultures incubated with either 4- or 5-cholestenone, or coprostanone that resulted in the production of coprostanol.^22^ This observation led us to propose that (*i*) the expression of *ismB* is not induced by cholesterol, (*ii*) there may be an isomerase enzyme not yet identified in *E. cop* that converts 4-cholestenone to 5-cholestenone, which *E. cop* IsmB then reduces to coprostanone, or (*iii*) there exist two 5β-reductases in *E. cop* that separately convert 4- and 5-cholestenone to coprostanone. Overall, these findings led us to propose a revised *ism* pathway in which the C3 hydroxyl in cholesterol is first oxidized by IsmA to produce 5-cholestenone (**Figure 4D**). The 5-cholestenone intermediate can be reduced by IsmB to produce coprostanone or can be non-productively isomerized to 4-cholestenone. A putative isomerase can convert 4-cholestenone back to 5-cholestenone. Another unknown 5β-reductase may catalyze the conversion of 4-cholestenone to coprostanone. Finally, the reduction of coprostanone by IsmA produces coprostanol.

### IsmA-encoding and IsmB-encoding human gut bacteria are associated with lower stool and serum cholesterol and elevated levels of stool 4-cholestenone and coprostanol

We next evaluated the extent to which the presence of IsmA-encoding and IsmB-encoding bacteria in complex gut microbial communities is associated with coprostanol production *in vivo*. We returned to the two independent cohorts with paired metagenomic and metabolomic data, HMP2 and PRISM, and categorized samples as being in groups that are categorized as having neither *ism* gene (*ismA*^-^/*ismB*^-^), only the *ismB* gene (*ismA*^-^/*ismB*^+^), or both *ism* genes (*ismA*^+^/*ismB*^+^) (**Figure 5A and 5B**). In both cohorts, samples from the *ismA*^+^/*ismB*^+^ group had significantly less (p-value ≦ 0.0001) cholesterol in their associated stool metabolomes compared to the *ismA*^-^/*ismB*^-^ and *ismA*^-^/*ismB*^+^ groups. Additionally, the *ismA*^+^/*ismB*^+^ group in both cohorts had significantly more 4-cholestenone and coprostanol in their stool metabolomes than the *ismA*^-^/*ismB*^-^ and *ismA*^-^/*ismB*^+^ groups. This supports the hypothesis that IsmA-encoding and IsmB-encoding bacteria within a gut microbiome convert intestinal cholesterol to coprostanol. Additionally, while the *ismB* gene in highly prevalent amongst human gut bacteria associated with stool coprostanol, in our analyses we find that it is the *ismA* gene that is necessary for this activity. Interestingly, we observed that in a subset of microbial communities from the *ismA*^-^/*ismB*^-^ and *ismA*^-^/*ismB*^+^ groups, coprostanol was still present in the associated stool metabolome. This observation could be explained by the limitations of metagenomic sequencing, especially with low abundance microbes, or the likelihood that distantly related bacteria and their 3β-hydroxysteroid dehydrogenases and 5β-reductases may perform the conversion of cholesterol to coprostanol.

**Figure 5.**
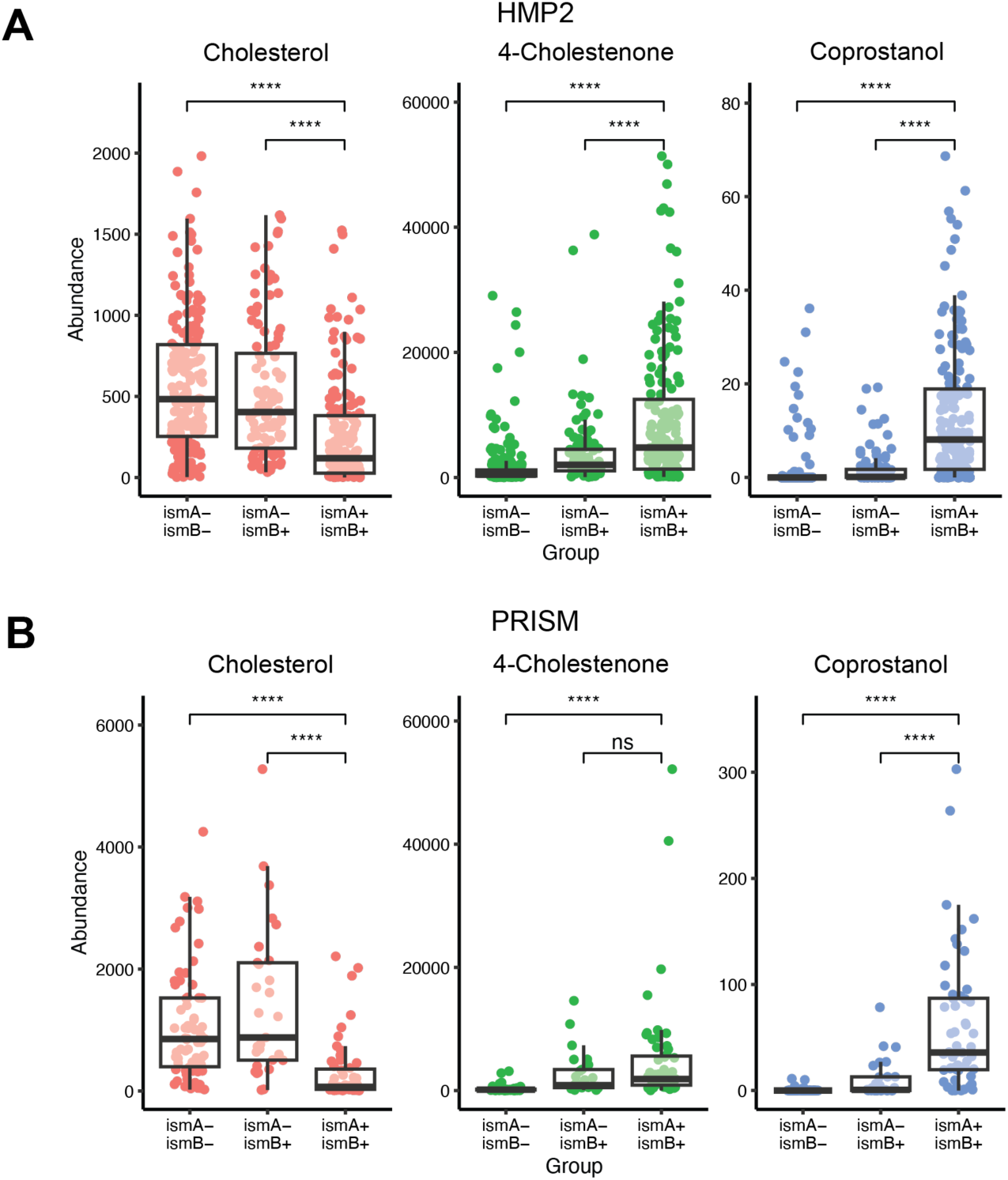
IsmA-encoding and IsmB-encoding human gut bacteria are associated with reduced stool cholesterol and greater stool coprostanol levels. Stool samples from human participants from the **(A)** HMP2 and **(B)** PRISM cohorts with *ismA*^+^/*ismB*^+^ MSPs have lower stool cholesterol and greater stool 4-cholestenone and coprostanol levels as determined by untargeted fecal metabolomics. Each point represents an independent sample, with the box representing the upper quartile, median, and lower quartile and the whiskers representing 1.5 times the interquartile range from the box (capped by the extrema of the data). Statistical significance and p-values were calculated using a two-sided Wilcoxon test; ns: p > 0.05; ****: p ≦ 0.0001.

We next evaluated the effect the presence of IsmA-encoding and IsmB-encoding bacteria may have on serum cholesterol levels in the FHS cohort. We found that the presence of *ismA* in human gut microbiomes is negatively associated with total cholesterol levels in serum metabolomes with a linear model corrected for age, sex, and BMI (standardized coefficient = −0.15, and 95% confidence interval = −0.24, −0.04]), consistent with our previous reported findings (**Table S1**).^12^ We found that the overwhelming majority of *ismA^+^* MSPs also encode *ismB* (18/19, 95%), and thus we cannot measure the effect of *ismA* alone on host serum cholesterol levels. Meanwhile, *ismB* alone does not show significant association with serum total cholesterol in a similar model (standardized coefficient = −0.03 and 95% confidence interval = [−0.18, 0.12]) (**Table S2**). These findings suggest that *ismA* is the critical gene that encodes the enzyme that potentially contributes to the decrease of host cholesterol levels, while the presence of *ismB* alone is not sufficient to have a significant effect on serum cholesterol levels. This work thus further supports our hypothesis that the conversion of cholesterol to coprostanol by *ismA*^+^/*ismB*^+^ gut microbiomes decreases host cholesterol levels.

## DISCUSSION

The notion that human gut bacteria metabolize intestinal cholesterol to coprostanol has existed for well over 100 years,^17,18^ but only recently have the genes and enzymes responsible for this conversion been investigated and biochemically validated.^12^ These efforts mark the start in obtaining a deeper understanding of the molecular determinants and the biological implications of this gut microbial activity. Related efforts in investigating the gut microbial transformation of cholesterol into cholesterol sulfate, and its role in modulating circulating levels of cholesterol in a gnotobiotic mouse model, exemplify the importance of linking gut bacteria-derived stool metabolites with enzymatic chemistry.^14,16^ The enzyme responsible for reducing a cholestenone intermediate to coprostanone was previously unknown, preventing the detection of the full *ism* metabolic pathway in gut microbial genomes and microbiomes as well as the genetic manipulation and/or engineering of the *ism* pathway. By combining bioinformatic searches, *in vitro* biochemical and culture-based assays, and human gut microbiome analyses, we identified and characterized IsmB, the gut bacterial enzyme responsible for the second step in coprostanol formation. IsmB catalyzes the reduction of 5-cholestenone, but not 4-cholestenone, into coprostanone under *in vitro* conditions, and its activity thus revises the previously proposed pathway for coprostanol production by gut bacteria. Previously, 5-cholestenone was not thought to be an on-pathway intermediate in coprostanol production, but rather was proposed to be a short-lived intermediate that is either enzymatically or non-enzymatically isomerized to 4-cholestenone, which is then converted to coprostanone.^22,60^ In our groups’ initial discovery of IsmA and its oxidative activity towards cholesterol, we overlooked the relevance of 5-cholestenone due to the isomerization of 5-cholestenone into 4-cholestenone under in vitro conditions and to the poor ionization of 5-cholestenone for LC–MS/MS quantitation, thus making it challenging to detect 5-cholestenone in our enzyme and lysate assays.^12^

Our newfound understanding of the specificity exhibited by IsmB for 5-cholestenone highlights a distinct mechanism by which Fe–S flavin-dependent 5β-reductases reduce non-conjugated β,γ-unsaturated steroidal substrates and sets precedence for this type of enzymatic transformation in the gut microbiome. It is intriguing that the gut microbial Fe–S flavoenzymes BaiCD^30^, Ci2350^37^, and OsrB^38^ all catalyze the reduction of steroidal substrates containing α,β-unsaturated bonds. While both Ci2350 and OsrB were found to have promiscuous activity towards a broad range of α,β-unsaturated steroid hormones^37,38^, the enzymes displayed either no or trace 5β-reductase activity toward 4-cholestenone, similar to *E. cop* IsmB. Even though 4-cholestenone is structurally similar to the steroid hormones metabolized by Ci2350 and OsrB, its non-polar lipid tail and different ᴅ-ring stereochemistry may prevent transformation by these 5β-reductases. This substrate preference is in contrast with the fact that conjugated α,β-unsaturated carbonyls are more electrophilic at the β position than unactivated alkenes, thus making them more susceptible to nucleophilic attack and reduction.^61^ The shared lack of 5β-reductase activity by Ci2350, OsrB, and *E. cop* IsmB toward 4-cholestenone further supports a distinct molecular mechanism for the gut bacterial conversion of cholesterol to coprostanol compared to other steroidal compounds that are metabolized by human gut bacteria, in which this transformation can proceeds through a 5-cholestenone intermediate. While the reduction of isolated alkenes by flavin-dependent enzymes has been reported,^62,63^ this is the first time, to the best of our knowledge, that this mode of reactivity has been observed for a human gut bacterial enzyme. The distinct substrate specificity exhibited by IsmB merits further investigation.

The biochemical specificity of IsmB for β,γ-unsaturated substrates has no precedence in naturally occurring enzymatic transformations of steroids. This is especially true for flavin-dependent OYE-like enzymes or enzymes with OYE domains which typically catalyze the reduction of olefins that are conjugated to polar groups, such as keto, carboxyl, cyano and nitro functional groups.^64^ The enzymatic function of IsmB merits further mechanistic study, as little is known about the substrate specificities of the larger class of microbial Fe–S flavin-dependent 5β-reductases, and how IsmB evolved to gain substrate specificity toward the β,γ-unsaturated 5-cholestenone intermediate. Furthermore, the large scale computational genomic analysis of Fe–S flavoenzymes from cultured and uncultured organisms completed by Andreu and coworkers^33^ revealed that this class of enzymes is predicted to be involved in the metabolism of a wide range of metabolites, including saccharides, amino acids, peptides, nucleotides, and lipids. Our phylogenetic analysis found that IsmB is part of this enzyme superfamily, highlighting the broad biochemical potential of that gut microbial Fe–S flavoenzymes, and represents a larger landscape of uncharacterized proteins that may mediate other important gut bacterial metabolic pathways, further supporting the need to biochemically investigate their functions.

Our work demonstrates that the majority of coprostanol-producing, IsmA-encoding and IsmB-encoding human gut bacteria form a distinct clade and phylogenetically separate themselves from IsmB-encoding bacteria that do not encode IsmA. The IsmB protein homologs identified in our phylogenetic analysis are encoded by predominantly biochemically uncharacterized and uncultured bacteria, exemplifying the challenges with investigating this metabolic pathway. We observed that uncultured gut bacterial species that carry both *ismA* and *ismB* are associated with lower levels of stool cholesterol, and greater levels of stool 4-cholestenone and coprostanol, further supporting our hypothesis that these bacteria may modulate host cholesterol levels. In order to accurately assess the associations between CVD risk and gut microbiome composition in future work, larger-scale prospective studies are required to explore this link as the cohorts used in this study are statistically underpowered.

One particularly challenging aspect of workflows for enzyme discovery is linking uncharacterized constitutively expressed genes to their biochemical functions.^65^ Unlike several other gut microbial reductive transformations of host derived metabolites or xenobiotics^66–69^, the expression of *E. cop* IsmB is not induced by cholesterol, and instead, IsmB appears to be constitutively expressed, and thus preventing us from searching gut bacterial transcriptomes to identify other human gut microbial IsmB homologs. Additionally, the *ismA* and *ismB* genes responsible for coprostanol formation in *E. cop* are not colocalized, unlike other gut bacterial metabolic pathways such as the *bai* operon responsible for intestinal bile acid dehydroxylation^27,30^, the SULT gene cluster for cholesterol sulfonation^14,16^, and the gut microbial pathway for the metabolism steroid hormones by *C. innocuum*.^37^ Gut microbial Fe–S flavin-dependent 5β-reductases are especially difficult to study because of the technical difficulties associated with expressing and purifying complex oxygen-sensitive metalloenzymes, and there are limited characterized examples from the human gut microbiome.^33^ We overcame these challenges by taking a chemically-guided genome mining approach to identify enzyme candidates for the conversion of 4-cholestenone to coprostanone and biochemically assessing their activities. Doing this revealed an unexpected substrate specificity for 5-cholestenone by IsmB. We applied this biochemical knowledge to search for and identify IsmB homologs in uncultured human gut bacteria. Additionally, paired analyses of human stool metagenomics and metabolomics helped support the functional role of IsmB in human gut bacterial conversion of cholesterol to coprostanol. Although the successful discovery and biochemical characterization of gut microbial enzymes enables assigning function to genes from cultured and uncultured human gut bacteria, the bioactivities and functions of microbial metabolites toward other gut microbes and the human host remain largely unknown.^1,4,70^ Our bioinformatic analyses of paired stool metagenomes and metabolomes revealed that individuals with *ismA*^+^/*ismB*^+^ gut microbiomes have decreased stool cholesterol and increased 4-cholestenone and coprostanol levels in their stool metabolomes. The activities of 4- and 5-cholestenone, coprostanone, and coprostanol all warrant further investigation, as the functions of these bacterial metabolites are not yet known. Other gut microbial metabolites that contain a core gonane skeleton, such as 3-oxolithocholic, isolithocholic acid and cholesterol sulfate, impact host biology and immune system regulation.^16,71,72^ It is this likely that 4- and 5-cholestenone, coprostanone, and coprostanol may also influence host metabolism and regulation, possibly through the activation of gut epithelial cell receptors. Our finding that the gut microbiome is correlated with lower host cholesterol levels^12^ and our discovery that IsmB can catalyze a plausible second step in the gut bacterial production of coprostanol will guide future experiments to investigate the potential causal role this activity in lowering cholesterol levels. Finally, this work highlights the importance of linking gut microbial biochemical activities to microorganisms, genes, and enzymes to fully understand metabolic interactions taking place within the human host.^68^

### Limitation of the study

We currently lack genetic tools needed to assess and validate the phenotypic function of *ismA* and *ismB* in *E. cop*, which could help further assign biochemical function to *ismA* and *ismB*. Not having access to genetically tractable *ismA*^+^/*ismB*^+^ coprostanol-producing gut bacterial strains prevents us from testing our central hypothesis that this gut microbial metabolic activity lowers host total cholesterol levels in an in vivo model. Efforts to culture and characterize gut Clostridial cholesterol-metabolizing bacteria lacking *ismA* or *ismB* will be necessary to further investigate the functional role of either *ism* genes in inducing a hypocholesterolemic host phenotype. Our ongoing search for a cholesterol transporter that may aid in the uptake of cholesterol, a potential isomerase that may catalyze the isomerization of 4-cholestenone back to 5-cholestenone, and an additional 5β-reductase that may catalyze the direct conversion of 4-cholestenone to coprostanone, respectively, impedes our identification of alternative pathways for coprostanol production and our complete understanding of the how gut bacterial conversion of cholesterol to coprostanol modulates host cholesterol levels. Lastly, most of the human participants in the FHS cohort dataset used in this study were not diagnosed with CVD at the time of sample collection. A follow up study on the same participants from the FHS cohort tracking progression of cardiometabolic status is currently underway.

## Supporting information

STAR Methods

---

**Figure S1.**
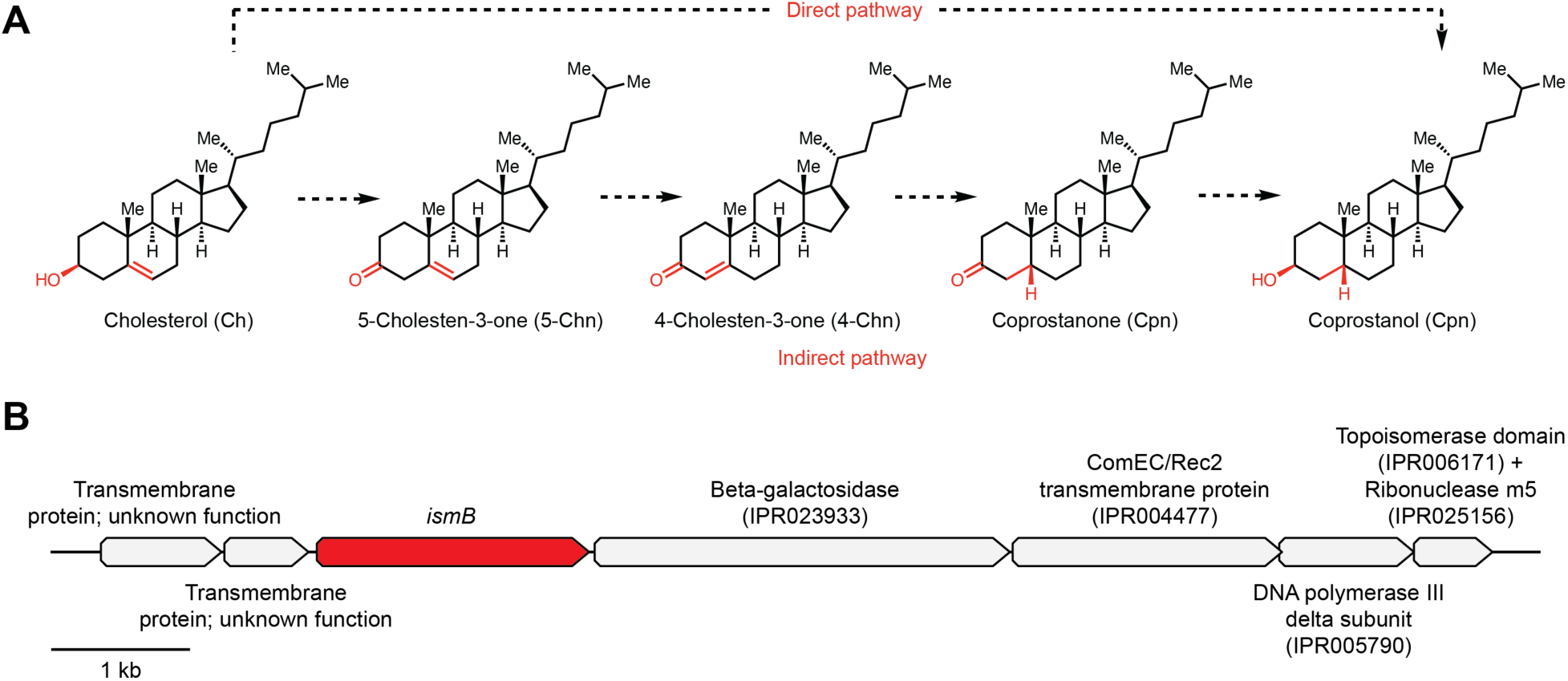
Direct and indirect pathways for the gut microbial metabolism of cholesterol to coprostanol and genome neighborhood of ECOP831. **(A)** Previously proposed direct and indirect pathways for the gut microbial conversion of cholesterol to coprostanol. The direct pathway implicates the conversion of cholesterol to coprostanol through the reduction of the C5–C6 double bond, in a single step, to the 5β-configured hydrogenated product coprostanol. The indirect pathway implicates a 3-step transformation involving the oxidation of cholesterol to 5-cholestenone, which then isomerizes into 4-cholestenone, followed by the reduction of the C4–C5 double bond to produce 5β-configured coprostanone, and a final reduction of the C3 keto group to the 3β-configured hydroxy group in coprostanol. **(B)** Genome neighborhood in *Eubacterium coprostanoligenes* ATCC 51222 (*E. cop*) containing ECOP831 and neighboring genes annotated with their predicted functions based on InterProScan.

**Figure S2.**
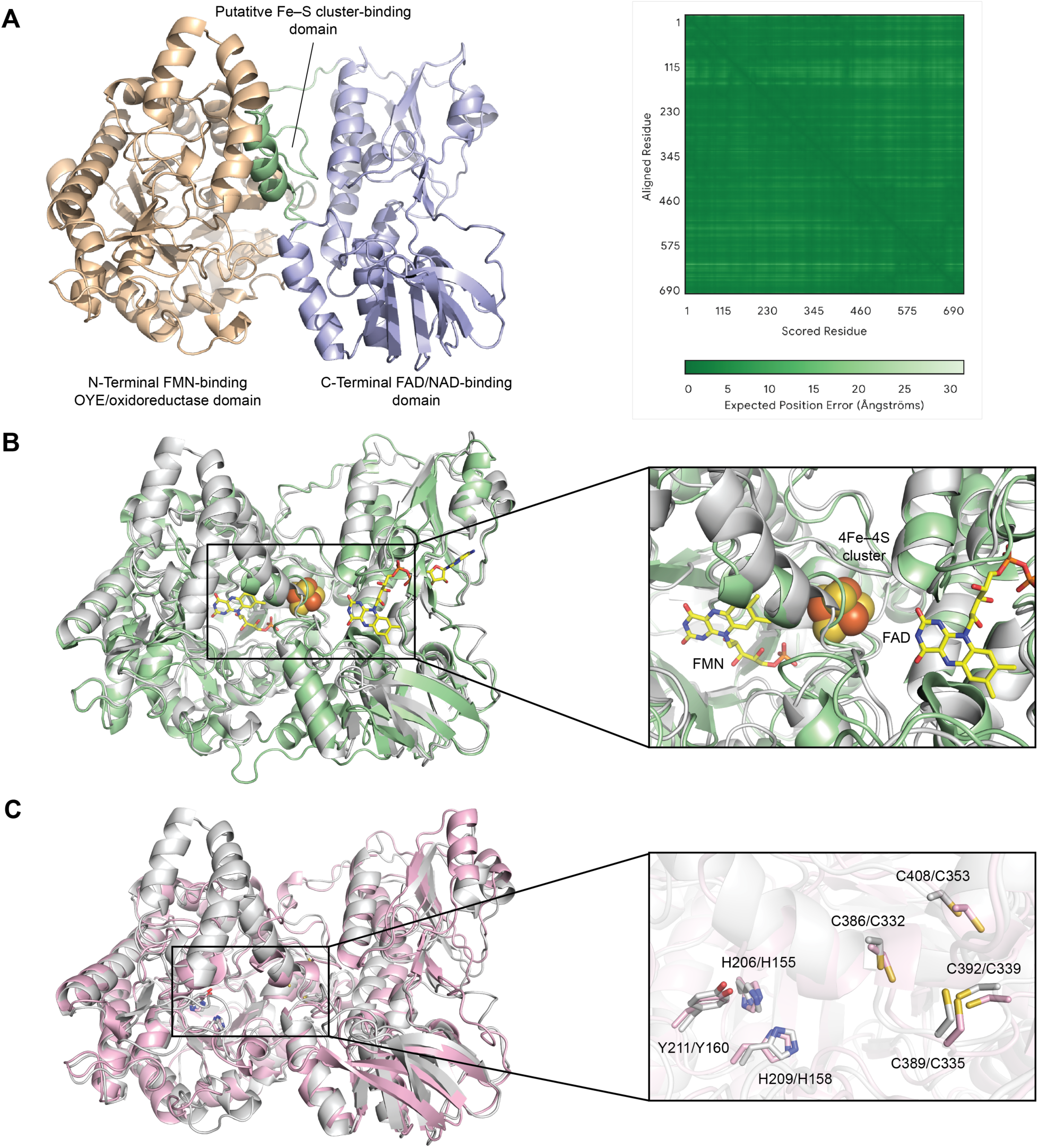
Predicted protein structure analysis of ECOP831 and other biochemically characterized Fe–S flavoenzymes. **(A)** AlphaFold3 predicted protein structure of ECOP831 obtained with high confidence scores and low predicted alignment errors. The N-terminal FMN-binding OYE/oxidoreductase domain is shown in wheat, the putative Fe–S cluster-binding domain is in pale green, and the C-terminal FAD/NAD-biding domain is in light blue. **(B)** Alignment of AlphaFold3 predicted structures of ECOP831 (gray) and the X-ray protein crystal structure of *E. coli* 2,4-dienoyl-CoA reductase (pale green) whose crystal structure contains protein-bound FMN and FAD cofactors, and a fully occupied [4Fe–4S] cluster, reveals a putative active site in the N-terminal domain for substrate binding and FMN cofactor binding, a centrally located cavity for the putative Fe–S cluster, and a C-terminal cavity for the binding of the FAD cofactor in ECOP831. **(C)** Alignment of AlphaFold3 predicted structures of ECOP831 (gray) and *C. scindens* BaiCD (pink) identifies Tyr and His residues hypothesized to be involved in substrate binding and catalysis in the OYE-like domain and cysteine amino acid residues in the ferredoxin-like domain for binding the putative Fe–S cluster cofactor.

**Figure S3.**
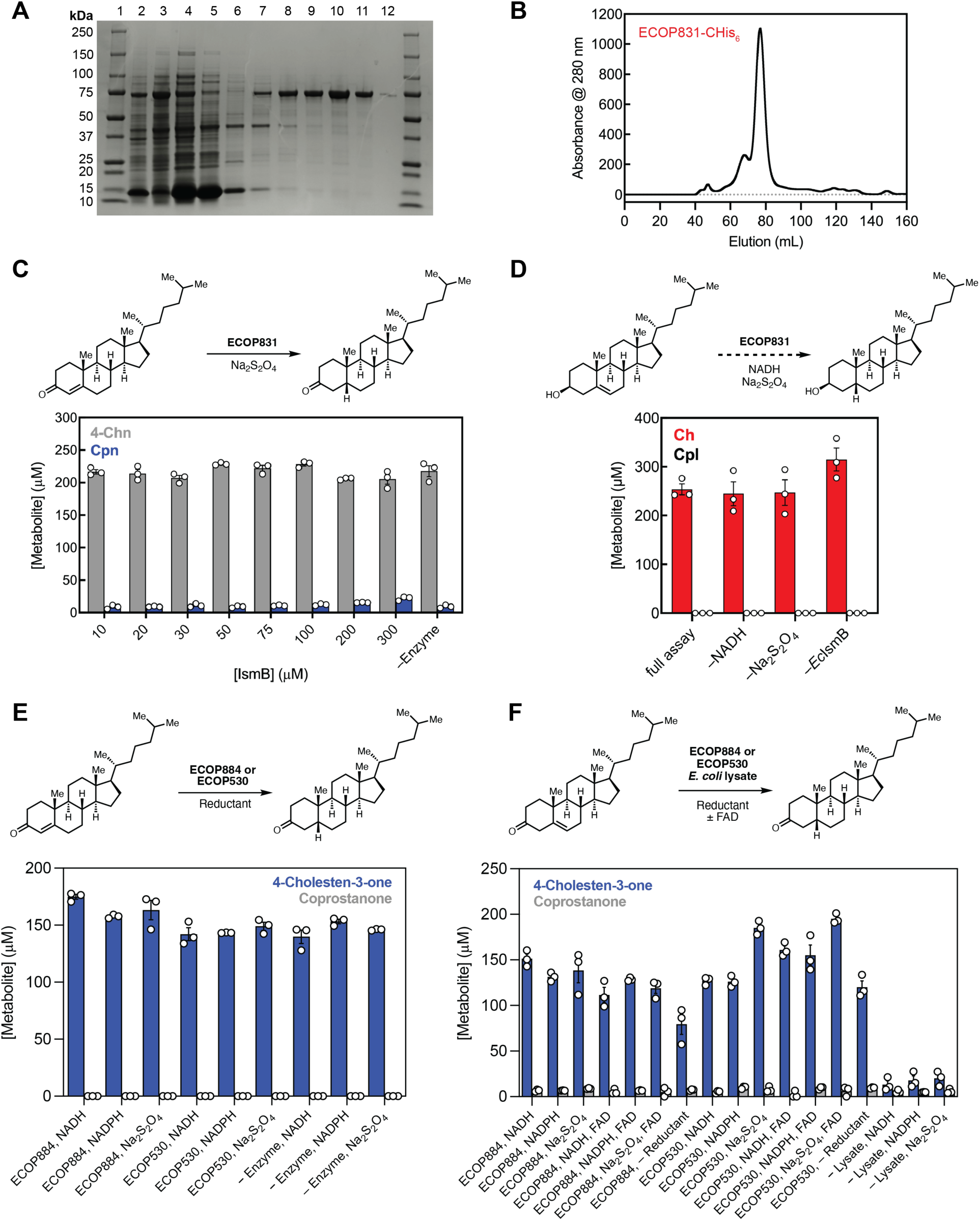
Characterization of ECOP831, ECOP884, and ECOP530 enzyme function. **(A)** SDS-PAGE of purified recombinant ECOP831 containing a C-terminal His_6_-tag. Precision Plus Protein All Blue Standards (BioRad) (lane 1), soluble lysate (lane 2), insoluble lysate (lane 3), flowthrough (lane 4), wash (lane 5), 25 mM imidazole (lane 6), 50 mM imidazole fractions 1 and 2 (lanes 7 and 8), 100 mM imidazole fractions 1 and 2 (lanes 9 and 10), and 300 mM imidazole fractions 1 and 2 (lanes 11 and 12). **(B)** Preparatory scale size exclusion chromatography of ECOP831-CHis_6_. **(C)** LC–MS/MS assays demonstrate ECOP831 does not efficiently convert 4-cholestenone into coprostanone. **(D)** LC–MS/MS assays with ECOP831as cholesterol as the substrate demonstrate that ECOP831 does directly reduce cholesterol to coprostanol. **(E)** LC–MS/MS assays with heterologously expressed and purified ECOP884 or ECOP530 using either NADH, NADPH, or Na_2_S_2_O_4_ as reductants demonstrate that these proteins do not convert 4-cholestenone to coprostanone. **(F)** LC–MS/MS assays with *E. coli* lysates heterologously expressing ECOP884 or ECOP530, using either NADH, NADPH, or Na_2_S_2_O_4_ as reductants, and with or without additional FAD cofactor demonstrate that these proteins do not convert 5-cholestenone to coprostanone. (C–F) n = 3 biologically independent replicates, data presented are the ± standard error mean (SEM).

**Figure S4.**
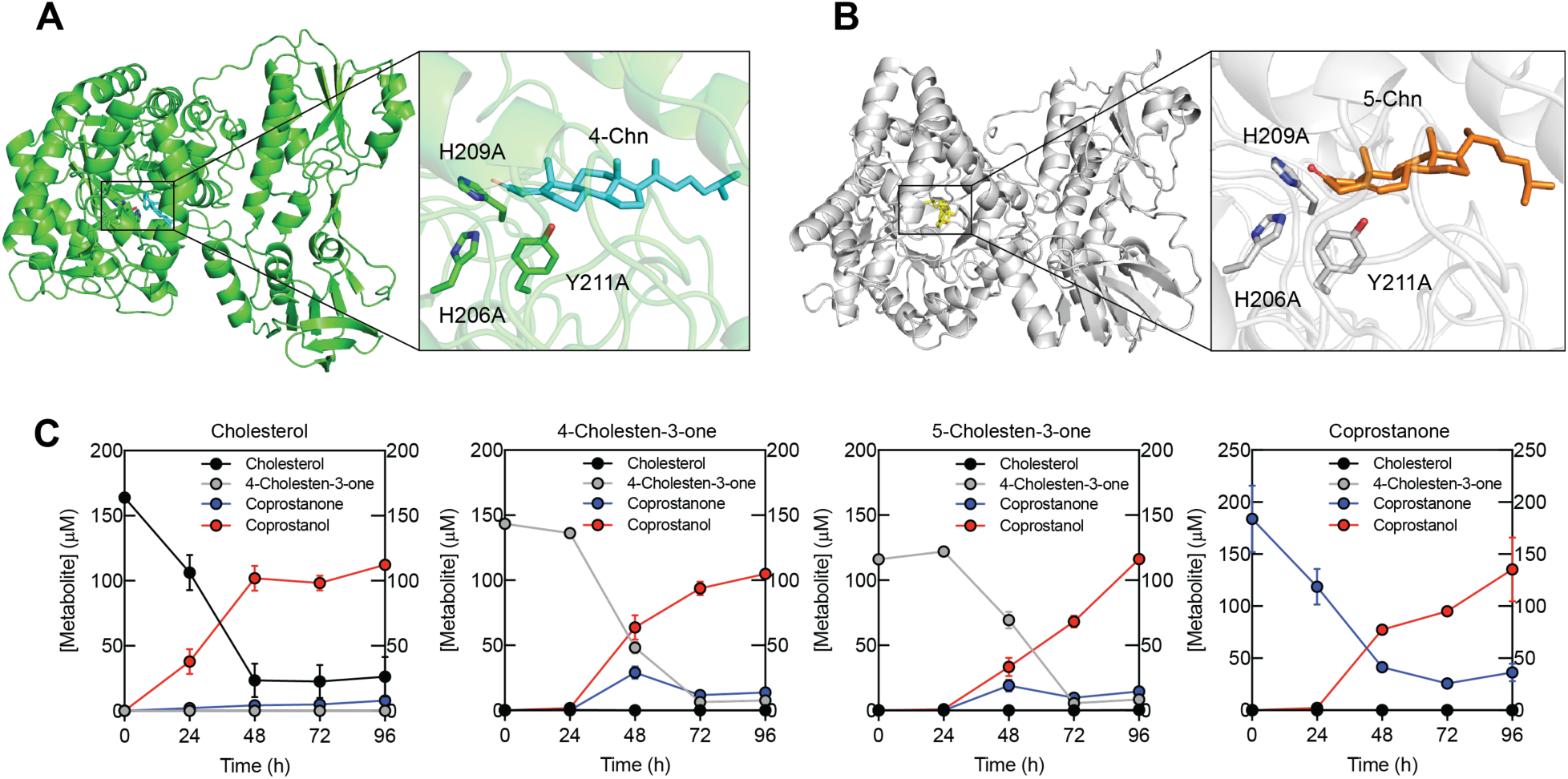
Substrate studies for *E. cop* IsmB and *E. cop* ATCC 51222. Docking models of the *E. cop* IsmB AlphaFold3 predicted structure with **(A)** 4-cholestenone and **(B)** 5-cholestenone using the molecular docking diffusion generative model DiffDock-L. **(C)** LC–MS/MS-based time course analysis of anaerobic *E. cop* liquid cultures for the conversion of cholesterol, 4-cholestenone, 5-cholestenone, and coprostanone to coprostanol. n = 3 biologically independent replicates, data presented are the ± standard error mean (SEM).

**Figure S5.**
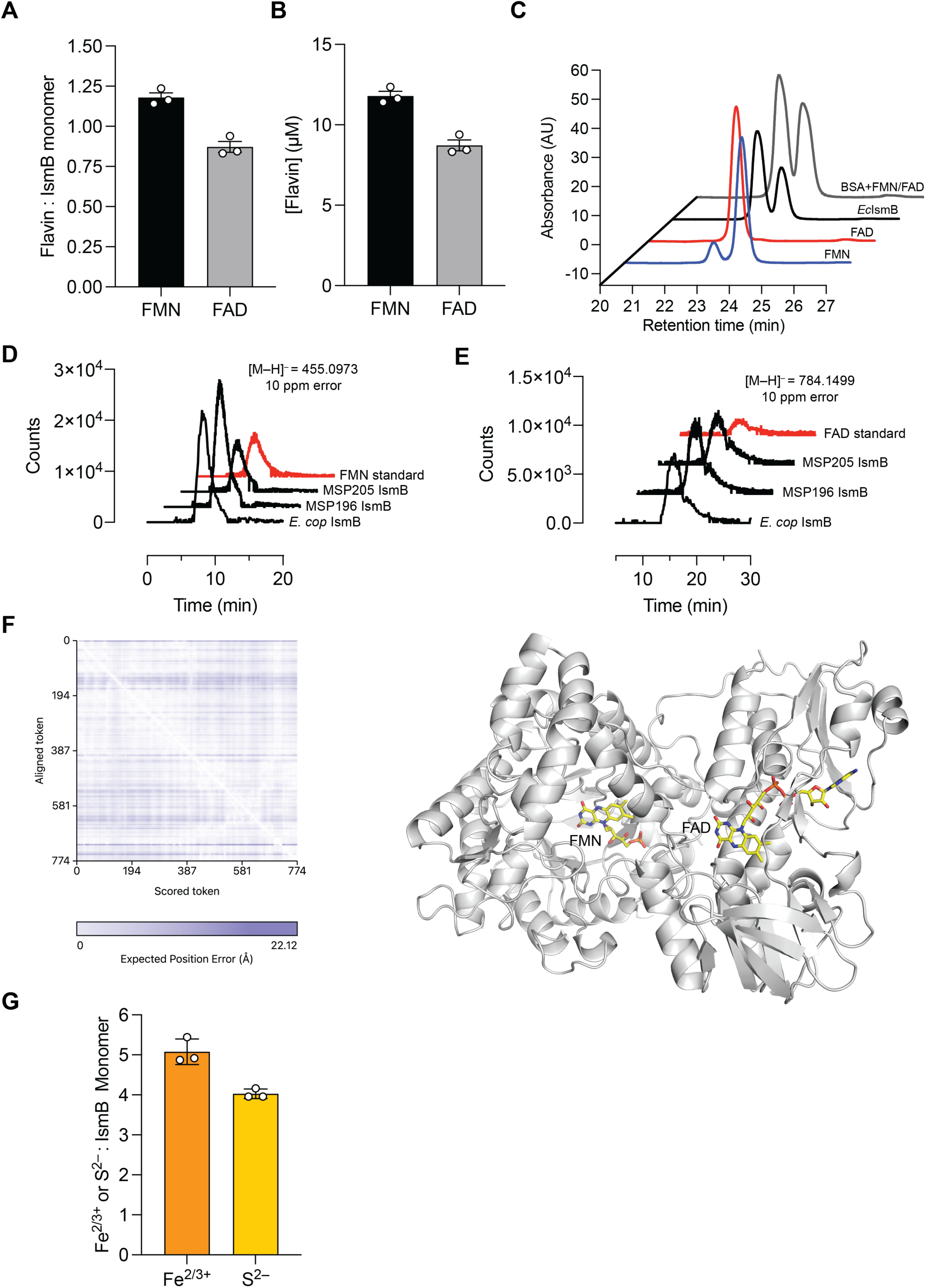
Cofactor binding analyses of *E. cop* IsmB. HPLC-based and LC–MS-based flavin quantitation assays, protein structure predictions, ICP–MS, and colorimetric assays reveal that *E. cop* IsmB binds both FMN and FAD, and a [4Fe–4S] cluster. **(A)** Flavin to IsmB monomer ratios and **(B)** FMN and FAD concentrations in 10 µM of *E. cop* IsmB. **(C)** HPLC–DAD traces of FMN and FAD standards, along with the extracted FMN and FAD cofactors found in *E. cop* IsmB and in BSA control spiked with FMN and FAD. High resolution quadrupole time-of-flight (Q-TOF) mass spectrometry LC–MS-based assays for the detection of **(D)** FMN and **(E)** FAD in IsmB homologs from *E. cop*, MSP196 and MSP205 reveal that all IsmB homologs bind both FMN and FAD. **(F)** Predicted protein structure of *E. cop* IsmB folded with FMN and FAD using Chai-1 reveals that the N-terminal OYE domain finds FMN while the C-terminal Rossman fold binds FAD. **(G)** Inductively-coupled plasma mass spectrometry-based (ICP–MS) iron quantitation and colorimetric sulfide content assays reveal that *E. cop* IsmB binds at least one [4Fe–4S] cluster. (A–E, G) n = 3 biologically independent replicates, data presented are the ± standard error mean (SEM).

**Figure S6.**
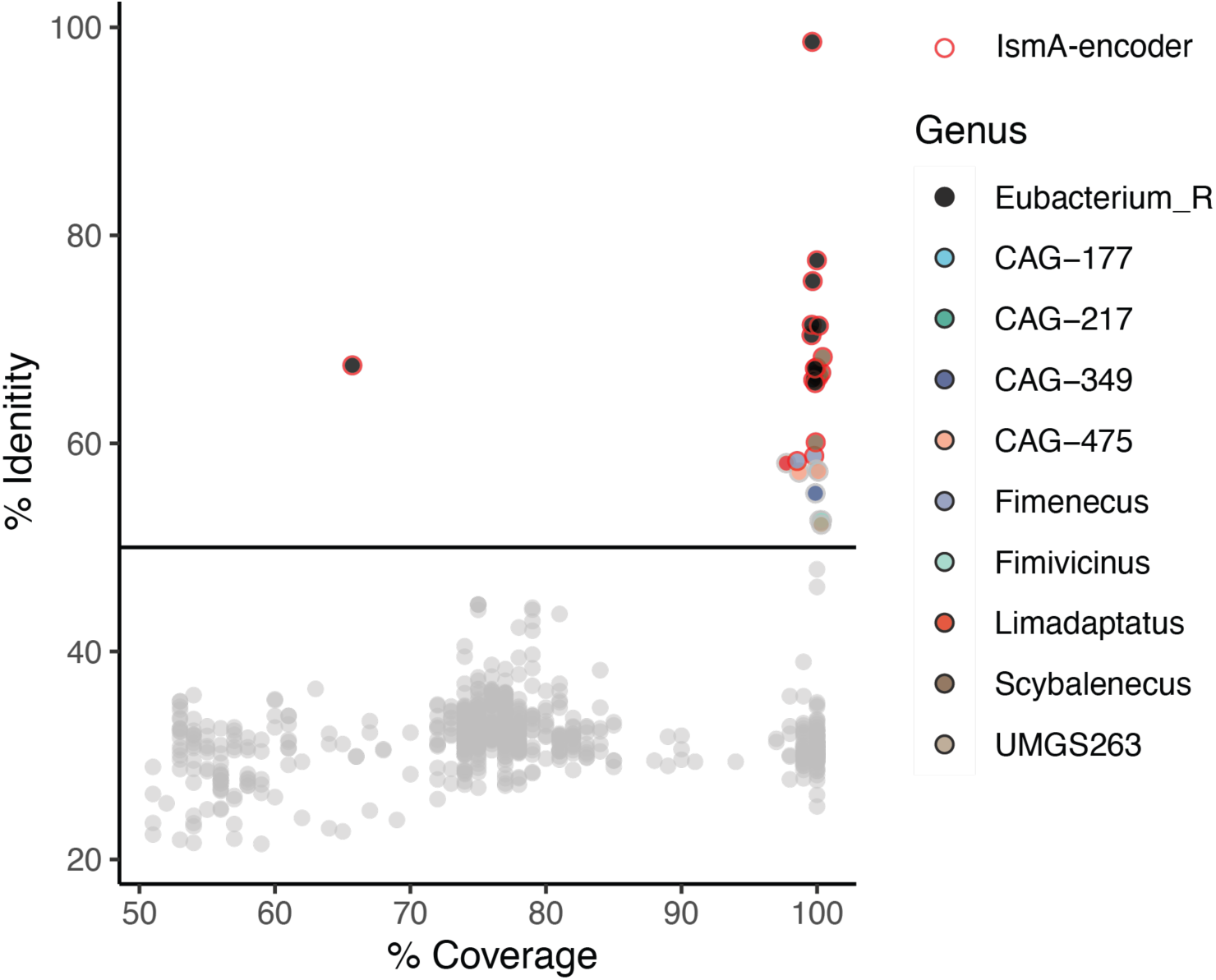
IsmB homologs encoded in uncultured human gut bacteria. **(A)** Jitter plot of 35 *E. cop* IsmB homologs (≥50% AA ID) found in human stool MSPs against percent protein sequence coverage. 19 MSPs were found to also encode IsmA homologs (red outline).

**Figure S7.**
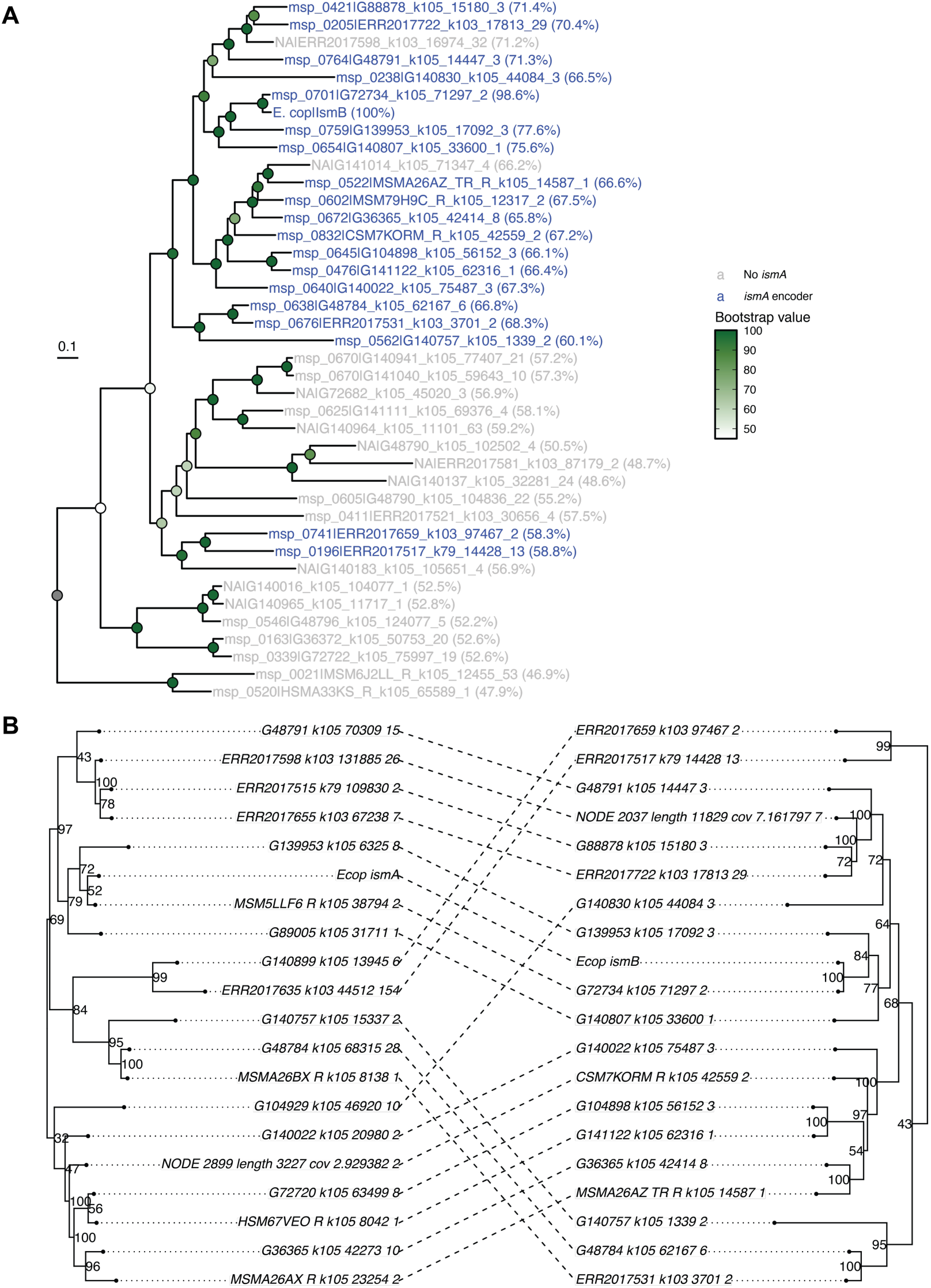
Protein phylogenetic analyses of human gut bacterial IsmB homologs. **(A)** Protein phylogenetic tree of IsmB homologs encoded by human gut MSPs and *E. cop* created using IQ-Tree. Homologs of IsmB colored in blue indicate that they are from bacteria or MSPs that encoded an *ismA* homolog, whereas gray-colored homologs are from MSPs that do not encode an *ismA* homolog. **(B)** Cophylogenetic analysis of IsmA and IsmB homologs. The low degree of entanglement in the phylogenetic tree suggests a low degree of incongruence between the evolution of IsmA and IsmB in coprostanol-producing bacteria.

**Figure S8.**
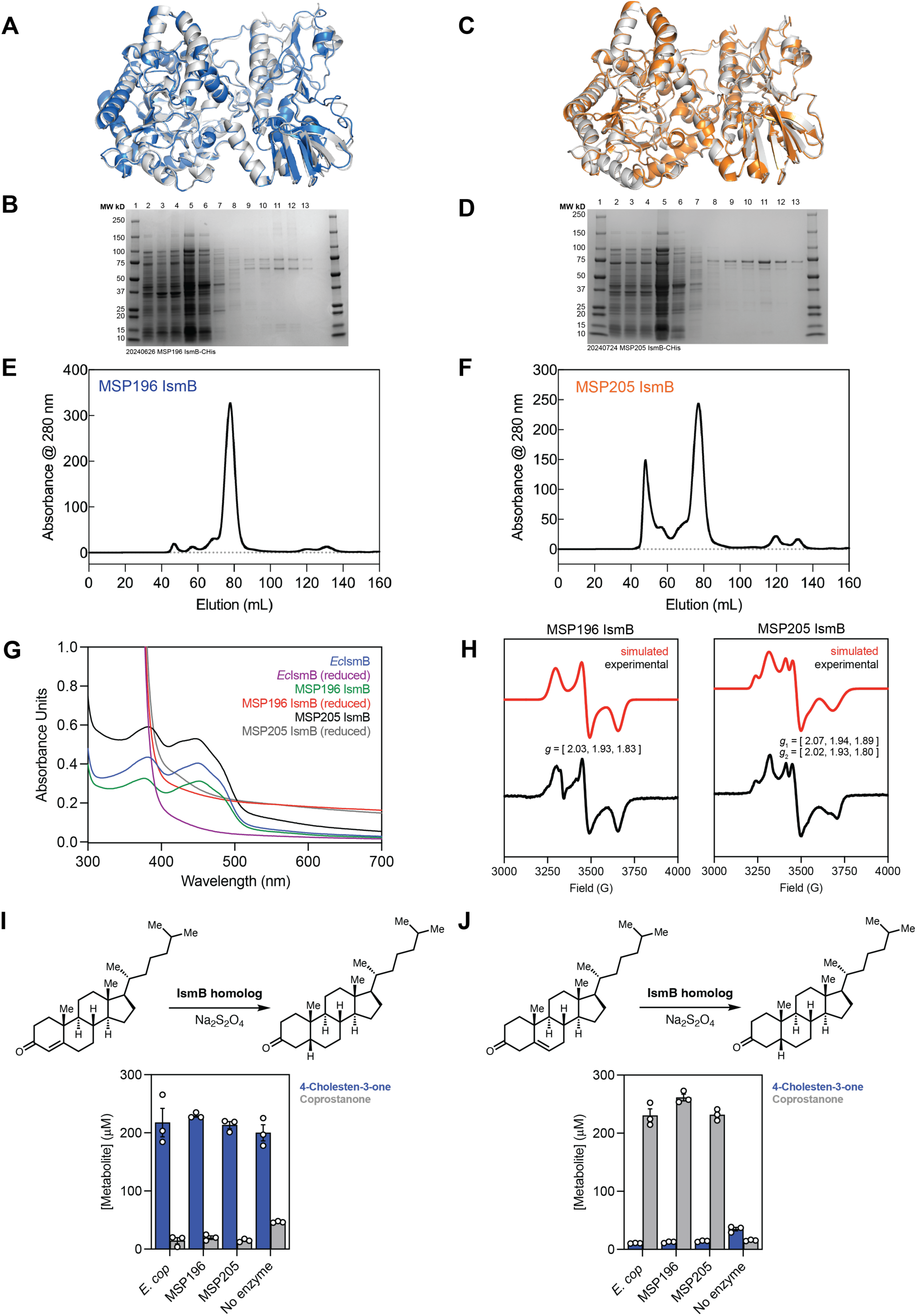
Characterization of human gut bacterial IsmB homologs. **(A)** AlphaFold3 predicted structure of MSP196 IsmB (blue) aligned with the AlphaFold3 predicted structure of *E. cop IsmB* (gray) with RMSD of 0.511 Å. **(B)** and **(D)** SDS-PAGE of purified recombinant IsmB homologs from MSP196 and MSP205 containing a C-terminal His_6_-tag. Precision Plus Protein All Blue Standards (BioRad) (lane 1), crude lysate (lane 2), soluble lysate (lane 3), insoluble lysate (lane 4), flowthrough (lane 5), wash (lane 6), 25 mM imidazole (lanes 6 and 7), 50 mM imidazole (lanes 8 and 9), 100 mM imidazole (lanes 10 and 11), and 250 mM imidazole (lanes 12 and 13). **(C)** AlphaFold3 predicted structure of MSP205 IsmB (orange) aligned with AlphaFold3 predicted structure of *E. cop* IsmB (gray) with RMSD of 0.326 Å. **(E** and **F)** Preparatory scale size exclusion chromatography of MSP196 IsmB-CHis_6_ and MSP205 IsmB-CHis_6_. **(G)** UV–vis spectroscopy of *E. cop* IsmB, and IsmB homologs from MSPs 196 and 205 supports that the human gut bacterial IsmB homologs also bind flavin cofactor(s) with absorption maxima at 380 and 450 nm. **(H)** EPR spectroscopy of IsmB homologs from MSPs 196 and 205 at 10 K with excess Na_2_S_2_O_4_ reveals rhombic spectra for both IsmB homologs, with the spectra of MSP 205 IsmB containing at least two different Fe–S cluster species, and indicates the presence of reduced [4Fe–4S] clusters. Anaerobic enzyme assays of IsmB homologs from *E. cop*, MSP 196, or MSP 205 using either **(I)** 4-cholestenone or **(J)** 5-cholestenone as the substrate indicate that IsmB from MSPs 196 and 205 also have substrate specificity for 5-cholestenone. (G–H) n = 3 biologically independent replicates, data presented are the ± standard error mean (SEM).

**Figure S9.**
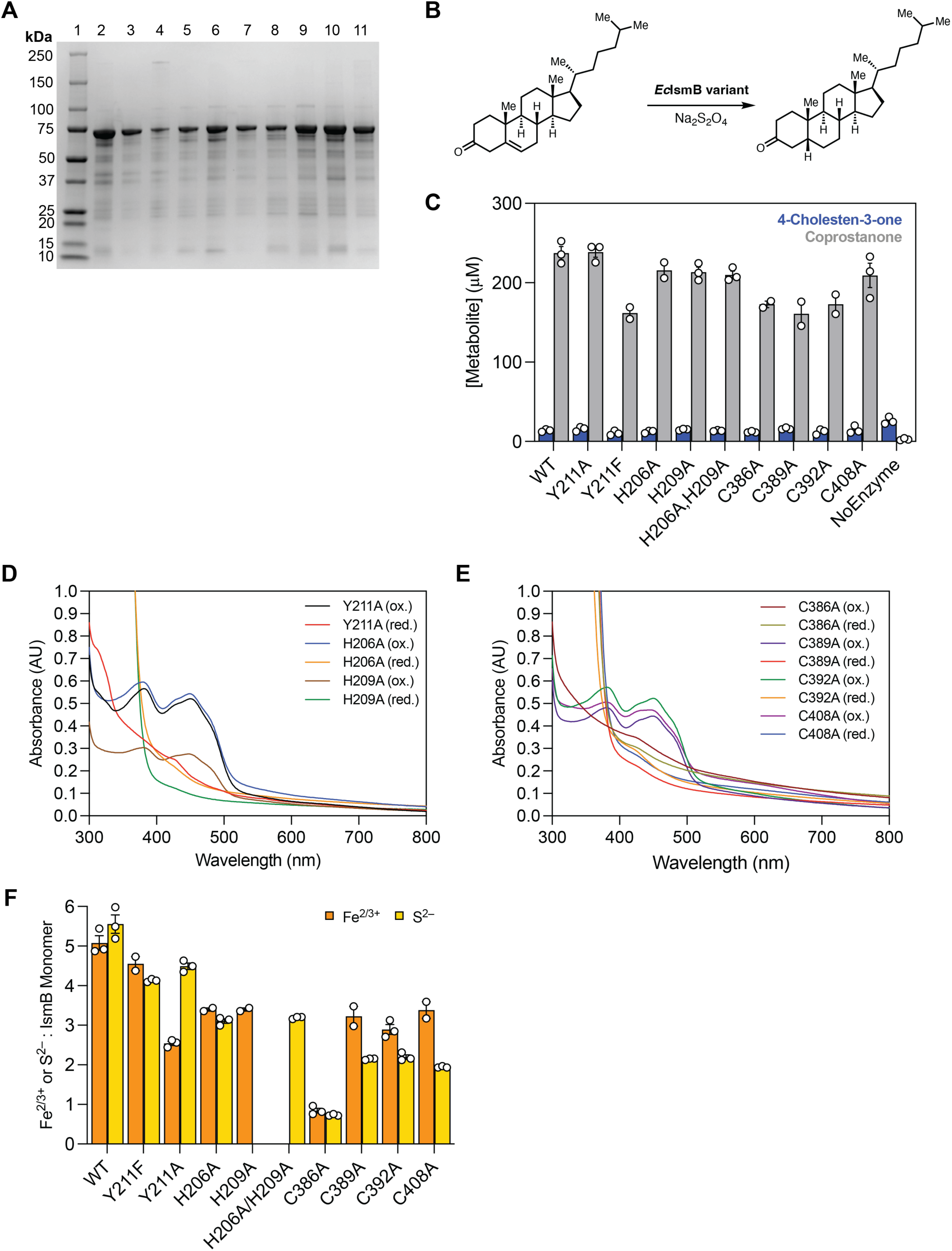
Characterization of *E. cop* IsmB active site variants. (**A)** SDS-PAGE of purified recombinant *E. cop* IsmB variants containing a C-terminal His_6_-tag. Precision Plus Protein All Blue Standards (BioRad) (lane 1), wild type (lane 2), Y211F (lane 3), C386A (lane 4), C389A (lane 5), C392A (lane 6), C408A (lane 7), H206A (lane 8), H209A (lane 9), Y211A (lane 10), and H206A/H209A (lane 11). **(B)** Chemical reaction catalyzed by *E. cop* IsmB active site variants. **(C)** LC–MS/MS assays with active site and cofactor binding site variants of *E. cop* IsmB. **(D)** UV–vis spectroscopy of *E. cop* IsmB active site variants Y211A, H206A, and H209A. **(E)** UV–vis spectroscopy of *E. cop* IsmB active site variants C386A, C389A, C392A, and C408A. **(F)** ICP–MS iron quantitation and colorimetric sulfide content assays reveal that *E. cop* IsmB active site variants have modulated Fe^2/3+^ and S^2–^ loading. (B, E) n = 3 biologically independent replicates, data presented are the ± standard error mean (SEM).

**Table S1.**
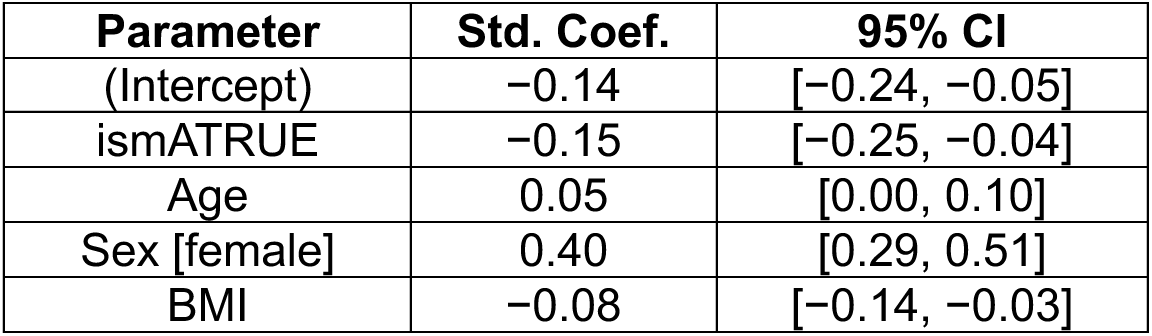
Association of total cholesterol levels with *ismA*^+^ encoder status in the FHS cohort using a linear model adjusting for age, sex and BMI (Std. Coef.: Standardized coefficient).

| Parameter | Std. Coef. | 95% CI |
| --- | --- | --- |
| (Intercept) | -0.14 | [-0.24, -0.05] |
| ismATrue | -0.15 | [-0.25, -0.04] |
| Age | 0.05 | [0.00, 0.10] |
| Sex [female] | 0.40 | [0.29, 0.51] |
| BMI | -0.08 | [-0.14, -0.03] |

**Table S2.** Association of total cholesterol levels with *ismB*^+^ encoder status in the FHS cohort using a linear model adjusting for age, sex and BMI (Std. Coef.: Standardized coefficient).

| Parameter | Std. Coef. | 95% CI |
| --- | --- | --- |
| (Intercept) | -0.20 | [-0.35, -0.05] |
| ismBTrue | -0.03 | [-0.18, 0.12] |
| Age | 0.05 | [0.00, 0.10] |
| Sex [female] | 0.40 | [0.29, 0.51] |
| BMI | -0.09 | [-0.14, -0.03] |

