## Supplementary material for "Discovery of the metalloenzyme IsmB revises a pathway for coprostanol formation by the human gut microbiome": STAR Methods

### KEY RESOURCES TABLE

| REAGENT or RESOURCE | SOURCE | IDENTIFIER |
|---|---|---|
| **Bacterial and virus strains** | | |
| *Eubacterium coprostanoligenes* ATCC 51222 | American Tissue Culture Collection | ATCC 51222 |
| *E. coli* NEB 5-alpha | New England Biolabs | C2987 |
| *E. coli* BL21(DE3) | New England Biolabs | C2527 |
| **Chemicals, Peptides and Recombinant Proteins** | | |
| β-Nicotinamide adenine dinucleotide hydrate | Millipore-Sigma | 43410 |
| β-NADH, disodium salt | Millipore | 481913 |
| Sodium hydrosulfite (sodium dithionite) | Sigma–Aldrich | 71699 |
| Cholesterol | Sigma–Aldrich | C8667 |
| 4-Cholesten-3-one | Cayman Chemical | 25713 |
| 5-Cholesten-3-one | Sigma–Aldrich | C75004 |
| Coprostanone | Steraloids Inc. | C5250-000 |
| Coprostanol | Cayman Chemical | 26764 |
| Cholesterol-25,26,26,26,27,27,27-$d_7$ | Avanti Polar Lipids | 700041 |
| L-α-Phosphatidylcholine (Lecithin) | Alfa Aeser | J61675-22 |
| Luria-Bertani Lennox (LB) medium | Research Products International | L24360 |
| Luria-Bertani Lennox (LB) medium, Agar | BD Difco | 240110 |
| Gifu anaerobic medium (GAM) | HiMedia | M1802 |
| 2XYT broth | Research Products International | X15600 |
| Isopropyl β-D-1-thiogalactopyranoside (IPTG) | Teknova | I3301 |
| Kanamycin sulfate | VWR | 97061-602 |
| Chloramphenicol | Sigma–Aldrich | C0378 |
| Riboflavin 5′-monophosphate sodium salt hydrate | Sigma–Aldrich | F6750 |
| Flavin adenine dinucleotide disodium salt | Cayman Chemical | 18167 |
| Phusion High-Fidelity DNA Polymerase | New England Biolabs | M0530S |
| Chicken egg lysozyme, salt free | Research Products International | L38100 |
| Pierce Protease inhibitor, EDTA-free | ThermoFisher Scientific | A32965 |
| InstantBlue Coomassie Protein Stain | Abcam | AB119211 |
| 2-mercaptoethanol | Sigma–Aldrich | 63689 |
| HEPES | Sigma–Aldrich | H4034 |
| Trizma hydrochloride | Sigma–Aldrich | T5941 |

| | | |
|---|---|---|
| Potassium chloride | Sigma–Aldrich | P9541 |
| Immidazole | ThermoFisher Scientific | A10221.22 |
| Glycerol | Sigma–Aldrich | G6279 |
| Deoxyribonuclease I from bovine pancreas | Sigma–Aldrich | DN25 |
| Magnesium chloride | Sigma–Aldrich | 208337 |
| Rubidium chloride | Sigma–Aldrich | 83979 |
| Nitric acid, trace metal | Supelco | NX0408 |
| Sodium sulfide nonahydrate | Sigma–Aldrich | 431648 |
| Iron (III) chloride | Sigma–Aldrich | 157740 |
| *N*,*N*-dimethyl-*p*-phenylenediamine | ThermoFisher Scientific | A15962.0B |
| **Oligonucleotides** | | |
| Primers for site directed mutagenesis of ECOP831 | Azenta | N/A |
| **Recombinant DNA** | | |
| Geneblocks ligated into pET28a/b vectors | Azenta, Genscript | N/A |
| pPH149-*iscSUA-hscBA-Fd* | Dr. John Latham; Latham et al. | N/A |
| **Deposited data** | | |
| *E. coprostanoligenes* ATCC 51222 genome | Kenny et al. | NCBI: PRJNA559861 |
| Framingham Heart Study | Kenny et al., Li et al. | NCBI: PRJNA559860<br>FHS repository: https://www.framinghamheartstudy.org/ fhs-for-researchers/researchapplication/ |
| Human Microbiome Project 2 (HMP2) | Lloyd-Price et al. | NCBI: PRJNA398089 |
| Prospective Registry in IBD Study at MGH (PRISM) | Franzosa et al. | NCBI: PRJNA400072 |
| **Software and algorithms** | | |
| GraphPad Prism 9 or 10 | GraphPad Software | N/A |
| EasySpin pepper (6.0.10) | EasySpin | https://easyspin.org/easyspin/documentation/index.html |
| MMseq2 | Steinegger et al. | https://github.com/soedinglab/MMseqs2 |
| MAFFT | Katoh | https://mafft.cbrc.jp/alignment/software/ |
| trimAl | Capella-Gutiérrez, S. et al. | https://github.com/inab/trimal |
| FastTree | Price at al. | https://github.com/morgannprice/fasttree |
| USEARCH | Edgar, R.C. et al. | https://github.com/rcedgar/usearch12 |
| GTDB-Tk | Chaumeil, P.A. et al. | https://github.com/Ecogenomics/GTDBTk |
| InterProScan | Jones et al. | https://www.ebi.ac.uk/interpro/search/sequence/ |
| DiffDock | Corso et al. | https://github.com/gcorso/DiffDock |
| IQ-TREE | Nguyen et al. | https://iqtree.github.io/ |
| AlphaFold3 | Abramson et al. | https://alphafoldserver.com |
| Chai-1 | Boitreaud et al. | https://github.com/chaidiscovery/ |
| PyMOL 2 | The PyMOL Molecular Graphics System, Version 3.0 | N/A |

### EXPERIMENTAL MODEL AND STUDY PARTICIPANT DETAILS

#### Microbial strains

*Eubacterium coprostanoligenes* (*E. cop*) ATCC 51222 was purchased from American Tissue Culture Collection. *E. cop* was cultured using Hungate tubes (Chemglass Life Sciences, catalog # CLS-4209-01) or using 50 mL conical tubes (Eppendorf, catalog # 0030122178) at 37 °C inside a vinyl anaerobic chamber (Coy Laboratory Products) under an atmosphere of <3.5% H2, 10% CO2, and N2 as the balance gas. *E. cop* was grown on Gifu Anaerobic Medium (HiMEDIA, catalog # M1801) supplemented with 1 g/L of soybean lecithin (Alfa Aeser, catalog # J61675-22). The GAM-lecithin broth was made anaerobic by autoclaving in 500 mL glass bottle, cycling the bottle into the vinyl anaerobic chamber while hot and allowing for gas exchange for a minimum of 24 h.

The strains *E. coli* NEB 5-alpha and BL21(DE3) were purchased from New England Biolabs, and were cultured on either sterile LB solid agar media or LB liquid media containing the appropriate antibiotics. To prepare chemically competent *E. coli* cells for plasmid transformations, the appropriate strain was cultured from a single colony in 100 mL of autoclaved 2XYT medium (Research Products International, Corp.) at 37 °C, and harvested at $OD_{600}$ of 0.3–0.4, and resuspended in 30 mL of sterile Transformation Buffer 1 (TFB1) (100 mM RbCl, 50 mM $MnCl_2$, 30 mM KOAc, 10 mM $CaCl_2$, 15% glycerol at pH 5.8) and kept on ice for 90 min. The *E. coli* cells were centrifuged for 5 min (4000 *g*, 4 °C), and the supernatant was discarded carefully. The cells were then resuspended in 4 mL of ice-cold Transformation Buffer 2 (TFB2) (10 mM MOPS, 10 mM $RbCl_2$, 75 mM $CaCl_2$, 15% glycerol, pH 6.8). 100 μL aliquots were flash frozen in liquid nitrogen and stored at −80 °C. Plasmid transformations were performed by adding one or more plasmid DNA (100–150 ng), 25 μL of thawed chemically competent *E. coli* cells in TFB2 buffer on ice for 15 min. The cells were heat shocked at 42 °C for 45 seconds, recovered in 300 μL of sterile LB broth, and incubated at 37 °C with shaking for 1 h before plating on LB agar containing appropriate antibiotic(s).

METHOD DETAILS

**Plasmid construction**

Plasmid construction was carried out using standard molecular cloning techniques and thermocycling was carried out in a Bio-Rad C100 thermal cycler. Descriptions of strains and plasmids are listed in Table S1AB. PCR amplifications were carried out with Phusion High-Fidelity polymerase (New England Biolabs), following manufacturer instructions, and using a Bio-Rad C100 thermal cycler. Primers (Azenta Life Sciences) are listed in Table S2B. Typical PCR experiments (total volume 50 μL) contained 10 ng of template, 0.5 μM forward primer, 0.5 μM reverse primer, optional 3% DMSO, and 2× Phusion-HF Master Mix (25 μL) (New England Biolabs). PCR parameters were the following: i) initial denaturation (98 °C for 30 s), 35 cycles of ii) denaturation (98 °C for 10 s), iii) annealing (55 °C for 30 s), iv) extension (72 °C for 90 s), and v) a final extension step (72 °C for 5 min). All PCR amplifications were analyzed by 1% agarose gel electrophoresis with ethidium bromide staining in 1 × Trisacetate-EDTA (TAE) buffer. Linearized plasmid vector was digested and gel purified, and both PCR products and linearized DNA were purified using Zymoclean DNA Clean and Concentrator Kit (Zymo Research). The circularized DNA constructs were assembled using Gibson Assembly (New England Biolabs), following manufacturer instructions. Constructs were verified by sanger sequencing (Azenta Life Science). Plasmid DNA was purified using Qiagen QIAprep Spin Miniprep Kit following manufacturer directions and transformed into chemically competent *E. coli* NEB 5-alpha cells (New England Biolabs) using protocol described above. Transformed cells were stored at −80 °C as frozen 25% glycerol stocks.

*Construction of plasmids for the heterologous expression of E. cop IsmB and IsmB homologs from human gut MSPs*. pET28b-EcopIsmB was constructed by ligating the synthetic gene insert *E. cop ismB* (UniProt ID: A0A1T4KLK7) containing the pET28b adapter sequences 'ForwardAdapter' and 'ReverseAdapter' onto a linearized pET28b backbone using Gibson Assembly. pET28b-MSP196IsmB and pET28b-MSP205IsmB were constructed similarly. All oligonucleotides used for plasmid construction are listed in Table S3.

*Site-directed mutagenesis of E. cop IsmB.* Amino acid residue variants of *E. cop* IsmB were prepared using a modified QuikChange mutagenesis protocol. Typical PCR experiments (total volume 25 μL) contained 2.5 ng of parental template DNA, 0.5 μM forward mutagenic primer, 0.5

μM reverse mutagenic primer, 3% DMSO, 2× Phusion-HF Master Mix (12.5 μL), and nuclease-free molecular biology grade water was added to make up the volume. PCR parameters were the following: i) initial denaturation (98 °C for 2 min), 25 cycles of ii) denaturation (98 °C for 20 s), iii) annealing (55–65 °C for 30 s), iv) extension (72 °C for 3.5 min), and v) a final extension (72 °C for 10 min). The PCR products were treated with Dpn1 restriction enzyme (New England Biolabs) at 37 °C for 1 hour to digest the template DNA, followed by heat inactivation at 80 °C for 20 min. To a 50 μL aliquot of chemically competent *E. coli* 5-alpha cells was added 10 μL of digested PCR product, and the transformed cells were plated onto $LB_{Km}$ agar plates. Single colonies were picked, grown out in liquid $LB_{Km}$ medium, and the plasmids were extracted using a QIAprep Spin Miniprep Kit (Qiagen). IsmB variants were verified by Sanger sequencing (Azenta Life Science) and whole plasmid Oxford Nanopore long read sequencing (Plasmidsaurus). All oligonucleotides used for mutagenesis are listed in Table S3.

**Expression and purification of heterologously expressed enzymes**

*Expression and purification of $His_6$-tagged IsmB enzymes*. *E. cop* IsmB, IsmB homologs from metagenomic species and active site variants were expressed and purified in a similar fashion. Chemically competent *E. coli* BL21(DE3) was transformed with the appropriate pET28b expression plasmid and the pPH149 plasmid encoding the *iscSUA-hscBA-Fd* genes for Fe–S cluster assembly (gifted by Dr. John A. Latham).[1] A 5 mL starter culture of the expression strain was grown in LB medium containing 50 and 34 μg $mL^{-1}$ kanamycin (Km) and chloramphenicol (Chl), respectively, by picking a single colony or scraping a glycerol stock. A 2.8 L anaerobic Erlenmeyer baffled flask containing 2 L of LB with 50 μg $mL^{-1}$ of Km and 34 μg $mL^{-1}$ Chl was supplemented with 200 μM $FeCl_3$ and 1 mM riboflavin 5′-monophosphate (flavin mononucleotide) and was inoculated with 5 mL of the starter culture. The culture was grown aerobically at 37 °C at 200 rpm until an $OD_{600}$ of 0.6–0.7 was reached. The cultures were chilled on ice for 15 min, induced with 500 μM IPTG, supplemented with 600 μM L-cysteine, tightly capped with a septum-lined twist cap, and grown anaerobically overnight at 20 °C at 180 rpm. Cell pellets were harvested by centrifugation at 10,000 × *g* for 15 min at 4 °C, flash frozen in liquid $N_2$, and stored at −80 °C.

All subsequent purification steps were performed in a hard shell anaerobic chamber at 4 °C containing 97% $N_2$ and 3% $H_2$ atmosphere (Coy Laboratory Products). *E. cop* IsmB was purified using buffers Ism Buffer 1 (B1) (50 mM HEPES, 300 mM KCl, 4 mM imidazole, 10 mM 2-mercaptoethanol, 10% glycerol, pH 7.5) and ism Buffer 2 (B2) (50 mM HEPES, 300 mM KCl, 500 mM imidazole, 10 mM 2-mercaptoethanol, 10% glycerol, pH 7.5). To a 50 mL conical tube containing the frozen cell pellet was added 10 mg $L^{-1}$ DNase I, 10 mM $MgCl_2 \cdot 6H_2O$, 1 mM flavin mononucleotide, 1 mM flavin adenine dinucleotide and a crushed Pierce™ Protease Inhibitor Tablet, EDTA-free (Thermo Scientific). The tube was cycled into the anaerobic chamber and the cell pellet was thawed, resuspended to approximately 5 mL of anaerobic Ism B1 per gram of wet cell paste, and the cells were lysed by sonicating with a half-inch horn at 25% amplitude for 5 min (2 s ON followed by 8 s OFF). The resulting suspension became dark yellow-brown during lysis. The lysate was transferred into a 50 mL high speed gas-tight centrifuge tube, taken out of the anaerobic chamber, and centrifuged at 20,000 × *g* for 30 min at 4 °C to separate soluble and insoluble fractions before being brought back into the anaerobic chamber. The soluble lysate loaded onto a Ni-NTA column (5 mL resuspended resin per 2 L expressed, pre-equilibrated with anaerobic ism B1 by gravity flow). The column was washed with 10 column volumes (cv) of 100% ism B1, then 90% ism B1/10% ism B2, 5 cv of 80% ism B1/20% ism B2, and 5 cv of 70% ism B1/30% ism B2. Protein was eluted from the column with 5 cv of 50% ism B1/50% ism B2 in 10 mL fractions. All fractions were analyzed via sodium dodecyl sulfate–polyacrylamide gel electrophoresis (SDS–PAGE) with Coomassie staining. The fractions containing purified IsmB were pooled and concentrated using an Ultra-15 Centrifugal Filters centrifugal concentrator with a 30 kDa MWCO membrane (Millipore Amicon) to a protein concentration of ~300 μM. The protein was buffer exchanged against ism Storage Buffer (SB) (20 mM HEPES, 300 mM KCl, 10 mM 2-mercaptoethanol, 10% glycerol, pH 7.5) three times. The protein concentration was measured in triplicate by method of Bradford using bovine serum albumin as a protein standard (Thermo Scientific). The protein solution was aliquoted into 0.5 mL amber-colored cryogenic tubes with an O-ring screw cap (Thermo Scientific). The tubes immediately frozen in liquid $N_2$, and stored at −80 °C.

*Determination of E. cop IsmB oligomeric state.* Anaerobic size-exclusion chromatography (SEC) was conducted inside an anaerobic chamber at 4 °C containing 97% $N_2$ and 3% $H_2$ (Coy Laboratory Products) with a ÄKTA Pure fast performance liquid chromatography system (Cytiva Life Sciences). *E. cop* IsmB was loaded onto a HiLoad 26/600 Superdex 200 pg size exclusion column (Cytiva Life Sciences) pre-equilibrated in ism SB. The resulting fractions were pooled, concentrated with a 30 kDa MWCO membrane (Millipore Amicon) and analyzed using SDS–PAGE with Coomassie staining.

*Expression and purification of His$_6$-tagged IsmA enzymes*. *E. cop* IsmA was expressed and purified under aerobic conditions. Chemically competent *E. coli* BL21(DE3) was transformed with the appropriate pET28b expression plasmid. A 5 mL starter culture of the expression strain was grown in LB containing 50 and 34 μg mL$^{-1}$ kanamycin (Km) by picking a single colony or scraping a glycerol stock. A 2.8 L Erlenmeyer baffled flask containing 2 L of LB with 50 μg mL$^{-1}$ of Km and was inoculated with 5 mL of the starter culture. The culture was grown aerobically at 37 °C at 200 rpm until an $OD_{600}$ of 0.6–0.7 was reached. The cultures were chilled on ice for 15 min, induced with 500 μM IPTG, and grown aerobically overnight at 20 °C at 180 rpm. Cell pellets were harvested by centrifugation at 10,000 × g for 15 min at 4 °C, flash frozen in liquid $N_2$, and stored at −80 °C.

All subsequent purification steps were performed under aerobic conditions at 4 °C. *E. cop* IsmA was purified using buffers ism Buffer 1 (B1) (50 mM HEPES, 300 mM KCl, 4 mM imidazole, 10 mM 2-mercaptoethanol, 10% glycerol, pH 7.5) and ism Buffer 2 (B2) (50 mM HEPES, 300 mM KCl, 500 mM imidazole, 10 mM 2-mercaptoethanol, 10% glycerol, pH 7.5). To a 50 mL conical tube containing the frozen cell pellet was added 10 mg L$^{-1}$ DNase I, 10 mM $MgCl_2 \cdot 6H_2O$, and a crushed Pierce™ Protease Inhibitor Tablet, EDTA-free (Thermo Scientific). The cell pellet was thawed, resuspended in approximately 5 mL per gram of wet cell paste with ism B1 and the cells were lysed by sonicating with a half-inch horn at 25% amplitude for 5 min (2.0 s ON followed by 8 s OFF). The resulting suspension became translucent off-white during lysis. The lysate was centrifuged at 20,000 × g for 30 min at 4 °C to separate soluble and insoluble fractions. The soluble lysate loaded onto a Ni-NTA column (5 mL resuspended resin per 2 L expressed, pre-equilibrated with anaerobic ism B1 by gravity flow). The column was washed with 10 column volumes (cv) of 100% ism B1, then 90% ism B1/10% ism B2, 5 cv of 80% ism B1/20% ism B2, and 5 cv of 70% ism B1/30% ism B2. Protein was eluted from the column with 5 cv of 50% ism B1/50% ism B2 in 10 mL fractions. All fractions were analyzed via sodium dodecyl sulfate–polyacrylamide gel electrophoresis (SDS–PAGE) with InstantBlue Coomassie staining (Abcam). The fractions containing purified IsmB were pooled and concentrated using an Ultra-15 Centrifugal Filters centrifugal concentrator with a 10 kDa MWCO membrane (Millipore Amicon) to a protein concentration of ~300 μM. The protein was buffer exchanged against ism Storage Buffer (SB) (20 mM HEPES, 300 mM KCl, 10 mM 2-mercaptoethanol, 10% glycerol, pH 7.5) three times. The protein concentration was measured in triplicate by method of Bradford using bovine serum albumin as a protein standard (Thermo Scientific). The protein solution was aliquoted into 1.5 mL microcentrifuge tubes and immediately frozen in liquid $N_2$, and stored at −80 °C.

*Expression and purification of His$_6$-tagged ECOP530 or ECOP884* BaiN homologs*.* ECOP530 and ECOP884 were expressed and purified under aerobic conditions. Chemically competent *E. coli* BL21(DE3) were transformed with the synthesized pET28a expression plasmid containing either the *ECOP530* or *ECOP884* gene with an N-terminal His$_6$ tag (GenScript). A 5 mL starter culture of the expression strain was grown in LB containing 50 and 34 µg mL$^{-1}$ kanamycin (Km) by picking a single colony or scraping a glycerol stock. A 2.8 L Erlenmeyer baffled flask containing 2 L of LB supplemented with 1 mM riboflavin 5′-monophosphate (flavin mononucleotide) and 50 µg mL$^{-1}$ of Km was inoculated with 5 mL of the starter culture. The culture was grown aerobically at 37 °C at 200 rpm until an $OD_{600}$ of 0.6–0.7 was reached. The cultures were chilled on ice for 15 min, induced with 500 µM IPTG, and grown aerobically overnight at 20 °C at 180 rpm. Cell pellets were harvested by centrifugation at 10,000 × g for 15 min at 4 °C, flash frozen in liquid $N_2$, and stored at −80 °C.

All subsequent purification steps were performed under aerobic conditions at 4 °C. ECOP530 and ECOP884 were purified using buffers Tris Buffer 1 (TB1) (20 mM Tris, 150 mM NaCl, 5 mM imidazole, 10 mM 2-mercaptoethanol, 10% glycerol, pH 7.0) and Tris Buffer 2 (TB2) (20 mM Tris, 150 mM NaCl, 500 mM imidazole, 10 mM 2-mercaptoethanol, 10% glycerol, pH 7.0). To a 50 mL conical tube containing the frozen cell pellet was added 10 mg L$^{-1}$ DNase I, 10 mM $MgCl_2 \cdot 6H_2O$, and a crushed Pierce™ Protease Inhibitor Tablet, EDTA-free (Thermo Scientific). The cell pellet was thawed, resuspended in approximately 5 mL per gram of wet cell paste with ism B1 and the cells were lysed by sonicating with a half-inch horn at 25% amplitude for 5 min (2 s ON followed by 8 s OFF). The resulting suspension became translucent and off-white during lysis. The lysate was centrifuged at 20,000 × g for 30 min at 4 °C to separate soluble and insoluble fractions. The soluble lysate loaded onto a Ni-NTA column (5 mL resuspended resin per 2 L expressed, pre-equilibrated with anaerobic ism B1 by gravity flow). The column was washed with 10 column volumes (cv) of 100% TB1, then 90% TB1/10% TB2, 5 cv of 80% TB1/20% TB2, and 5 cv of 70% TB1/30% TB2. Protein was eluted from the column with 5 cv of 50% TB1/50% TB2 in 10 mL fractions. All fractions were analyzed via sodium dodecyl sulfate–polyacrylamide gel electrophoresis (SDS–PAGE) with InstantBlue Coomassie staining (Abcam). The fractions containing purified ECOP530 or ECOP884 were pooled and concentrated using an Ultra-15 Centrifugal Filters centrifugal concentrator with a 10 kDa MWCO membrane (Millipore Amicon) to a protein concentration of ~300 µM. The protein was buffer exchanged against Tris Storage Buffer (TSB) (20 mM Tris, 150 mM NaCl, 10 mM 2-mercaptoethanol, 10% glycerol, pH 7.0) three times. The protein concentration was measured in triplicate by method of Bradford using bovine serum albumin as a protein standard (Thermo Scientific). The protein solution was aliquoted into 1.5 mL microcentrifuge tubes and immediately frozen in liquid $N_2$, and stored at −80 °C.

**End-point enzyme activity assays for detecting cholesterol metabolite production**

*IsmB anaerobic enzyme assays.* Enzyme activity assays were prepared by combining 3 mM $Na_2S_2O_4$ and 20 µM *E. cop* IsmB or IsmB variant in anaerobic Ism Reaction Buffer (RB) (50 mM HEPES, 50 mM KCl, pH 7.5) in a 1.5 mL microcentrifuge tube inside an anaerobic chamber containing $N_2$, and < 0.1 ppm $O_2$ at 22 °C (MBraun). The samples were thoroughly mixed prior to the addition of 300 µM of either cholesterol, 4-cholesten-3-one, 5-cholesten-3-one (5.0 mM stock solution in MeOH). After incubating at room temperature for 18 h, the reaction mixtures were taken out of the anaerobic chamber and quenched with 9 equivalents of cold MeOH containing 11.1 µM cholesterol-d$_7$ as an internal standard. The samples were vortexed at high speed for 2 min, centrifuged at 16,000 rpm for 10 min at 4 °C, and then transferred to a screw top propylene 250 µL LC–MS vial (Agilent).

*IsmA aerobic enzyme assays.* Enzyme activity assays were prepared by combining 20 μM *E. cop* IsmA and either 3.0 mM β-nicotinamide adenine dinucleotide phosphate ($NADP^+$) or β-nicotinamide-adenine dinucleotide phosphate, reduced (NADPH) in aerobic Ism Reaction Buffer (RB) (50 mM HEPES, 50 mM KCl, pH 7.5) in a 1.5 mL microcentrifuge tube. The samples were thoroughly mixed prior to the addition of either 300 μM cholesterol or 5-cholesten-3-one (5.0 mM stock solution in MeOH). After incubating at room temperature for 16 h, the reaction mixtures were quenched with 9 equivalents of cold MeOH containing 11.1 μM cholesterol-$d_7$ as an internal standard. The samples were vortexed at high speed for 2 min, centrifuged at 16,000 rpm for 10 min at 4 °C, and then transferred to a screw top propylene 250 μL LC–MS vial (Agilent).

*Aerobic enzymes assays with ECOP530 or ECOP884.* Enzyme activity assays were prepared by combining 20 μM *ECOP530 or ECOP884,* 3.0 mM of either β-nicotinamide-adenine dinucleotide, reduced (NADH) or β-nicotinamide-adenine dinucleotide phosphate, reduced (NADPH), and 20 μM of FAD in a 1.5 mL microcentrifuge tube containing aerobic Tris Base Reaction Buffer (20 mM HEPES, 150 mM NaCl, pH 7.0). The samples were thoroughly mixed prior to the addition of either 300 μM 4-cholesten-3-one or 5-cholesten-3-one (5.0 mM stock solution in MeOH). After incubating at 37 °C for 16 h, the reaction mixtures were quenched with 9 equivalents of cold MeOH containing 11.1 μM cholesterol-$d_7$ as an internal standard. The samples were vortexed at high speed for 2 min, centrifuged at 16,000 rpm for 10 min at 4 °C, and then transferred to a screw top propylene 250 μL LC–MS vial (Agilent).

**UPLC–MS/MS Methods**

Simultaneous analysis of cholesterol, 4-cholesten-3-one, 5-cholesten-3-one, coprostanone, and coprostanol was carried out by ultra performance liquid chromatography tandem mass spectrometry (UPLC–MS/MS). Liquid chromatography was performed using a Waters Acquity UPLC H-Class System (Waters Corporation). In a typical run, 1 μL of each sample was injected onto an Acquity UPLC BEH C8 1.7 μm (2.1 × 50 mm) column (Waters Corporation). The flow rate was 0.5 mL $min^{-1}$ using mobile phase A = 0.1% formic acid in $H_2O$ and mobile phase B = 0.1% formic acid and 5 mM ammonium acetate in MeOH. The column temperature was kept at 40 °C. An isocratic mobile phase of 10% A and 90% B was applied for 5 min. The first 0.5 min of the run was diverted to waste. MS detection was performed with a Waters Xevo TQ-S (Waters Corporation) instrument with atmospheric pressure chemical ionization in positive mode (APCI+) (capillary voltage, 3.10 kV; cone voltage, 42 V; source offset voltage, 50 V; desolvation temperature, 500 °C; desolvation gas flow, 800 L $h^{-1}$; cone gas flow, 150 L $h^{-1}$; nebulizer, 7.0 bar). Tandem MS/MS analysis parameters were optimized using authentic standards, unless otherwise stated. See Table S4 for specific detection parameters for precursor and daughter ions.

**Characterization by electron paramagnetic resonance spectroscopy (EPR)**

Wild-type *E. cop* IsmB was prepared for EPR spectroscopy as follows. All assay concentrations are final concentrations. Anoxic *E. cop* IsmB aliquots were brought into an anaerobic chamber containing $N_2$ and < 0.1 ppm $O_2$ at 22 °C (MBraun). A 200 mM sodium hydrosulfite ($Na_2S_2O_4$) solution was prepared in anaerobic ism Reaction Buffer (RB) (50 mM HEPES, 50 mM KCl, pH 7.5). To a 1.5 mL microcentrifuge tube was added 200 μM *E. cop* IsmB and 15 mM $Na_2S_2O_4$ to a final volume of 250 mL, and the mixture was gently pipetted up and down resulting in a clear faint red protein solution. The entire 250 μL reaction mixture was used for analysis. The sample was loaded into an EPR tube with 4 mm outer diameter and 8″ length (Wilmad LabGlass, 734-LPV-7), sealed with a rubber NMR tube septum, and immediately flash frozen in liquid $N_2$.

All EPR experiments were conducted at either the MIT Department of Chemistry Instrument Facility (DCIF) or the Harvard University Laukien-Purcell Instrumentation Center. A Bruker EMX-Plus spectrometer with an ER4119HS high sensitivity X-band resonator in perpendicular mode in conjunction with a Bruker/ColdEdge 4K waveguide cryogen-free cryostat were used to carry the experiments at liquid helium temperatures. A Xenon 1.1b.155 software package was used for data acquisition. The magnetic field was calibrated with an external standard of $K_2(SO_3)_2NO$ (Frémy) [ $g_x$ = 2.00785, $g_y$ = 2.00590, $g_z$ = 2.00265, $A_x$ = 5.5 G, $A_y$= 5.0 G, $A_z$ = 28.7 G]. The experimental spectra were modeled with EasySpin (Version 6.0.10) for MATLAB (MathWorks) to obtain *g*-values, hyperfine coupling constants, and line widths.

EPR spectra were recorded under the following conditions: temperature, 10 K; center field, 3350 G; sweep width, 200 G; microwave power, 1.262 μW; microwave frequency, 9.37 GHz; modulation amplitude, 10 G; modulation frequency, 100 kHz; time constant, 0.01 ms; conversion time, 10.72 ms; sweep time, 16.08 s; receiver gain, 10 dB. Simulated spectra were integrated twice to quantify the number of spins in each sample.

**Iron, sulfide and flavin detection, quantification and UV–vis assays**

The iron content of *E. cop* IsmB and IsmB variants was measured by inductively coupled plasma mass spectrometry (ICP–MS) at the MIT Department of Chemistry Instrument Facility. Briefly, *E. cop* IsmB was diluted to 1.5 mg $mL^{-1}$ in molecular biology grade water in a 1.5 mL microcentrifuge tube to a final volume of 200 μL. Then, 60 μL of trace-metal grade nitric acid (Millipore Sigma) was added, and the acidified protein solution was vortexed gently, and incubated at 60 °C for 3 hours to ensure complete denaturation of the protein. The protein samples were centrifuged at 16,000 rpm for 2 min, and the supernatants were transferred to a metal-free 15 mL conical tube (VWR) and diluted in molecular biology grade water up to 300 μL. The samples were measured using an Agilent 7900 ICP–MS instrument. Standards for $^{56}Fe$ were prepared from 1000 ppm Fe standard solution (SPEX Certiprep).

The sulfide content of a 10 μM solution of *E. cop* IsmB was determined using a modified published procedure, with the only differences being that assays were performed in microcentrifuge tubes, assay volumes were doubled, pipetting was used for mixing instead of stirring, and the mixture was incubated for 20 min after the addition of NaOH.

The identities of flavin cofactors bound to *E. cop* IsmB were assessed by reversed phase HPLC. Briefly, SEC-purified *E. cop* IsmB was diluted in triplicate in molecular biology grade water to 10 μM in a PCR tube strip to final volume of 300 μL. The protein samples were denatured by heating at 95 °C for 10 minutes using a Bio-Rad C100 thermal cycler, followed by centrifugation at 3,217 × *g* for 20 min. Avoiding the insoluble protein pellet, 200 μL of the supernatant was carefully transferred, filtered using a 0.2 μm Millex hydrophilic PTFE syringe filter (Millipore) to an amber-colored HPLC vial containing a spring-loaded glass insert (VWR). The samples were analyzed by HPLC using a Luna 5 μm C18(2) 100 Å LC Column (125 × 4 mm) (Phenomenex). 20 μL of each sample were injected onto the column using a Thermo Fisher Dionex-Ultimate3000 UHPLC connected to a diode array detector (DAD). The flow rate was 1 mL $min^{-1}$ using 5 mM sodium formate in water as mobile phase A and MeOH as mobile phase B. The column was maintained at 25 °C. The following gradient was applied: 0–5 min, 15% B isocratic; 5–30 min, 15–75% B; 30–35 min, 75–100% B; 35–50 min, 100–75% B. FMN and FAD were detected by measuring the absorbance at 264 nm. For quantification, FMN and FAD standards were prepared in sterile filtered $H_2O$ and a standard curve was measured in singlicates.

LC–MS detection of flavin cofactors was performed using a Q-TOF instrument. SEC-purified *E. cop* IsmB, MSP 196 IsmB and MSP 205 IsmB were diluted to 5 µM (275 µL final volume) in technical triplicates using degassed anaerobic miliQ water inside a glove box (MBraun) in 1.5 mL screw-cap microcentrifuge tubes. The samples were boiled at 99 °C for 10 min, and centrifuged at 16,000 rpm for 10 min at 4 °C. The supernatant was filtered through a low protein-binding hydrophilic LCR PTFE 0.2 µm syringe filter, and the filtrate was transferred into amber vials with a spring-loaded glass insert. 5 µL of each sample was analyzed by LC–MS using an Agilent Q-TOF 6530 equipped with a Dual AJS ESI source in negative mode and a Cogent 4 Diamond Hydride (4 µm, 100 Å, 150 × 3 mm) HPLC column heated to 30°C, at a flow rate of 0.5 mL/min. Solution A was $H_2O$ + 0.1% formic acid, and Solution B was acetonitrile + 0.1% formic acid. The LC method was as follows: 90% solution B for 1 min, 90–30% solution B for 19 min, 30% solution B for 1 min, 30–90% solution B over 4 min, hold at 90% solution B for 10 min. The following parameters were used for the Q-TOF: gas temp 300 °C, drying gas 11 L/min, nebulizer 35 psi, sheath gas temp 275 °C, sheath gas flow 11 L/min, VCap 3500 V, and nozzle voltage 500 V. Extracted ion chromatograms were obtained for FMN ($[M-H]^-$ = 455.0973 *m/z* and FAD ($[M-H]^-$ = 784.1499 *m/z*) and for representative replicates of each sample and 1 µM standard solutions of FAD, FMN or FAD + FMN. The EICs were plotted on GraphPad Prism (v. 9.3.1).

To obtain the UV–vis spectra of as isolated protein, *E. cop* IsmB was diluted to 20 µM with anoxic buffer (50 mM HEPES, 50 mM KCl, pH 7.5) and was transferred to an Ultra-Micro Cell quartz cuvette (Hellma) which was capped with a rubber septum inside of an anaerobic chamber 97% $N_2$ and 3% $H_2$ (CoyLabs). The absorbance of the solution was measured from 220 nm to 900 nm using a Cary 8454 UV-Vis Diode Array System (Agilent). To obtain a spectrum for the reduced protein, 20 mM sodium dithionite was added with a gas-tight syringe before the absorbance was measured at 5 min.

**Activity assays in *E. cop* Lysates**

*E. cop* was cultured anaerobically at 37 °C in 40 mL of GAM-lecithin broth, taking 1 mL of culture to measure growth using $OD_{600}$ measurements. When the cultures reached an $OD_{600}$ of 0.5 they were spiked with either 4 mL of 5.0 mM cholesterol in GAM-lecithin broth or with GAM-lecithin broth as the vehicle control to an approximate final concentration of 500 µM cholesterol and final volume of 40 mL of *E. cop* liquid culture. The cultures were incubated at 37 °C for 12 h, at which point they were harvested by centrifugation at 1,500 × *g* for 20 min at 4 °C, and the supernatant was discarded. In an anaerobic chamber under an atmosphere of < 3.5% $H_2$, 10% $CO_2$, and $N_2$ as the balance gas, the *E. cop* cell pellets were thawed and resuspended in 5 mL of ism RB containing 10 mM of 2-mercaptoethanol. The resuspended cells were lysed by sonicating with a half-inch horn at 25% amplitude for 2.5 min (2 s ON followed by 8 s OFF). The crude lysates were then incubated at room temperature for 18 h with 300 µM cholesterol, 4-cholesten-3-one or 5-cholesten-3-one and 3.0 mM $NADP^+$ or $Na_2S_2O_4$ in a final volume of 50 µL in 1.5 mL microcentrifuge tubes. Reaction mixtures were analyzed by LC–MS as described above using three different methods for detecting cholesterol, 4-cholesten-3-one and coprostanone.

**Phylogenetic analysis of bacterial Fe–S flavoenzyme superfamily**

The maximum-likelihood phylogenetic tree of Fe-S flavoenzyme superfamily proteins were created using a workflow adapted from a previous study.[2] Proteins sequences containing the two Pfam domains, Oxidored_FMN (PF00724) and Pyr_redox_2 (PF07992) were retrieved from UniprotKB (accessed July 07, 2025; N=25,695). Sequences with length between 600 and 800 amino acids were clustered using MMSeq2 (90% identity, 90% coverage).[3] The corresponding Pfam domains were extracted from the cluster representative sequences (N=10,840) with hmmsearch (v3.4) against PF00724 and PF07992 (sequences with length not shorter than 250 or 150 were kept respectively; N=9,440). Sequences were aligned using MAFFT (v7.453)[4] and the multiple sequence alignments (MSAs) were trimmed using trimAl (v1.5; -gt 0.9).[5] The trimmed MSAs of the two domains were merged and a phylogenetic tree was inferred using FastTree using default parameters (v2.1.10).[6]

**Metagenomics and metabolomics data analysis**

Metagenomic gene catalog creation and untargeted metabolomic data collection and analysis from fecal samples were performed on which datasets? using a previously published workflow.[7] The gut microbial IsmB proteins were identified from protein families of clustered sequences in the gene catalog (50% identity and 50% coverage) using usearch (v8.1)[8]. Metagenomic Species Pangenomes (MSPs) with the *ismB* gene were noted as the IsmB-encoding MSPs, and phylogenetic analysis was performed on the pangenomes of such MSPs and reference species (*E. coprostanoligenes* and *C. scindens*) with GTDB-TK (the de novo workflow; "--taxa_filter s__Eubacterium_R coprostanoligenes,s__Clostridium_AP scindens --outgroup_taxon s__Clostridium_AP scindens"; GTDB-Tk v2.4.0; GTDB v220)[9]. IsmA and IsmB sequences identified from the MSPs were aligned using MAFFT (v7.453)[4], and phylogenetic trees with bootstrap values were created with IQ-TREE (v2.3.6; "-B 1000")[10] for the multiple sequence alignments of IsmA and IsmB sequences, respectively.

**CITED WORK**
